# Structural basis of polyamine transport by human ATP13A2 (PARK9)

**DOI:** 10.1101/2021.05.28.446245

**Authors:** Sue Im Sim, Sören von Bülow, Gerhard Hummer, Eunyong Park

## Abstract

Polyamines are small, organic polycations that are ubiquitous and essential to all forms of life. Currently, how polyamines are transported across membranes is not understood. Recent studies have suggested that ATP13A2 and its close homologs, collectively known as P5B-ATPases, are polyamine transporters at endo-/lysosomes. Loss-of-function mutations of ATP13A2 in humans cause hereditary early-onset Parkinson’s disease. To understand the polyamine transport mechanism of ATP13A2, we determined high-resolution cryo-EM structures of human ATP13A2 in five distinct conformational intermediates, which together represent a near-complete transport cycle of ATP13A2. The structural basis of the polyamine specificity was revealed by an endogenous polyamine molecule bound to a narrow, elongated cavity within the transmembrane domain. The structures show an atypical transport path for a water-soluble substrate, where polyamines may exit within the cytosolic leaflet of the membrane. Our study provides important mechanistic insights into polyamine transport and a framework to understand functions and mechanisms of P5B-ATPases.

**Highlights:** Cryo-EM structures of human ATP13A2 in five distinct conformations at 2.5–3.7 Å resolutions.

Unique features of ATP13A2 in comparison to other P-type ATPases.

Structure of the substrate-binding pocket of ATP13A2 and the molecular basis of polyamine binding.

Conformational changes along the transport cycle and proposed model for polyamine transport.

## Introduction

P-type ATPases form a large superfamily of active transporters that catalyze directional translocation of substrates across cellular membranes (Clausen et al., 2017; Dyla et al., 2019; Kuhlbrandt, 2004; López-Marqués et al., 2020; Palmgren and Nissen, 2011). Most P-type ATPases function as either ion pumps (P1 to P3-ATPases) (Morth et al., 2007; Olesen et al., 2007; Shinoda et al., 2009; Sorensen et al., 2004; Toyoshima et al., 2000; Toyoshima et al., 2004) or lipid flippases (P4-ATPases) (Bai et al., 2019; Bai et al., 2020; Hiraizumi et al., 2019; Nakanishi et al., 2020; Timcenko et al., 2019). These transporters perform vital physiological tasks, such as generation of ionic gradients across membranes and lipid asymmetry in membrane leaflets.

Recent studies suggest that P5-ATPases, arguably the least characterized subfamily, have substrate specificities and functions that are critical for intracellular organelle homeostasis but fundamentally differ from those of P1-to P4-ATPases. Among the two subtypes—P5A and P5B (Møller et al., 2008; Sørensen et al., 2010), the P5A-ATPase (ATP13A1 in humans) seems to act as a protein quality control factor at the endoplasmic reticulum (ER), which extracts mis-targeted amino (N)-or carboxy (C)-terminally anchored transmembrane proteins from the ER membrane (Feng et al., 2020; McKenna et al., 2020; Qin et al., 2020). P5B-ATPases reside at endo-/lysosomal membranes (Sorensen et al., 2018), and humans express four isoforms, ATP13A2 to ATP13A5. Recent functional studies have suggested that ATP13A2, a prototypical P5B-ATPase, is a polyamine transporter at late endosomes and lysosomes (De La Hera et al., 2013; van Veen et al., 2020).

Polyamines are polycationic, aliphatic molecules that are ubiquitous and essential to all forms of life (Handa et al., 2018; Igarashi and Kashiwagi, 2010; Pegg, 2009). By binding to nucleic acids, proteins, and lipids, polyamines participate in numerous cellular processes, including modulation of chromatin structure, synthesis of proteins and nucleic acids, oxidative stress response, and ion channel functions (Handa et al., 2018; Pegg, 2009). The main naturally occurring forms are spermidine (triamine), spermine (tetraamine), and their precursor putrescine (diamine). In mammals, P5B-ATPases seem to play a key role in the uptake of extracellular polyamines by exporting polyamines from endocytic vesicles to the cytosol (De La Hera et al., 2013; Hamouda et al., 2020; Heinick et al., 2010; van Veen et al., 2020).

Defects in ATP13A2 are strongly implicated in progressive neurological disorders, such as Parkinson’s disease and hereditary spastic paraplegia (Di Fonzo et al., 2007; Estrada-Cuzcano et al., 2017; Ramirez et al., 2006; Yang and Xu, 2014). In ATP13A2 knock-out cells, lysosomes abnormally accumulate polyamines, resulting in lysosomal alkalization, dysfunction, and rupture (Dehay et al., 2012; van Veen et al., 2020). ATP13A2 deficiency also sensitizes cells to oxidative stress, suggesting a role of ATP13A2 in protecting cells against mitochondrial reactive oxygen species (Vrijsen et al., 2020). The involvement of ATP13A2 in maintaining lysosomal and mitochondrial health may explain the link between ATP13A2 defects and the pathogenesis of the diseases (Exner et al., 2012; Usenovic et al., 2012; Wallings et al., 2019). It has been suggested that small molecules that potentiate the polyamine transport activity of ATP13A2 might enhance endo-/lysosomal functions and provide therapeutic effects for neuroprotective treatments (Gitler et al., 2009; van Veen et al., 2020). However, the mechanistic basis of polyamine transport by ATP13A2 remains to be elucidated.

In the present study, we determined high-resolution cryo-electron microscopy (cryo-EM) structures of human ATP13A2 in five distinct conformations along its polyamine transport cycle. The chemical structure of polyamines is markedly different from any other characterized substrates of P-type ATPases due to their elongated structure and regularly spaced positive charges. Our study elucidated how ATP13A2 has adapted to specifically recognize these chemical signatures of polyamines and enable their membrane translocation. We also revealed how the common architecture of P5-ATPases could give rise to the strikingly different substrate specificities (polypeptides vs. polyamines) between the P5A and P5B subtypes. Together with mutagenesis analysis and molecular dynamics simulations, our work offers mechanistic insights into polyamine transport and provides a framework for further functional investigations and pharmacological targeting of P5B-ATPases.

## Results

### Cryo-EM analysis of human ATP13A2

To determine the structure of ATP13A2, we expressed full-length human ATP13A2 in *Spodoptera frugiperda* (Sf9) cells using baculovirus. ATP13A2 was extracted from membranes with a mixture of dodecyl maltoside (DDM) detergent and cholesteryl hemisuccinate (CHS) and isolated to apparent homogeneity by affinity purification with a green fluorescent protein (GFP) tag attached to the C-terminus of ATP13A2. Purified ATP13A2 eluted largely as a single peak in size-exclusion chromatography and appeared as well-dispersed particles in cryo-EM images (Fig. S1 A–D).

The catalytic cycle of P-type ATPases involves ATP binding, phosphorylation of a conserved aspartate (D508 in ATP13A2), and subsequent dephosphorylation, processes which are mediated by the cytosolic domains of the transporter. Specifically, P-type ATPases follow a sequence of intermediate states, E1 (apo) → E1-ATP → E1P-ADP → E1P → E2P → E2-Pi → E1, in a scheme known as the Post-Albers cycle, where E1 and E2 represent two main conformations of the enzyme (Dyla et al., 2019; Kuhlbrandt, 2004; Palmgren and Nissen, 2011) (Fig. S1E). These transitions accompany conformational changes in the transmembrane domain (TMD) of the enzyme, which drive substrate transport. To stabilize the intermediate states of ATP13A2, we prepared wild-type (WT) and mutant enzymes (defective in phosphorylation or substrate binding) in the presence of an adenine nucleotide (ADP, ATP, or β,γ-methyleneadenosine 5’-triphosphate [AMP-PCP]) or phosphate analog (BeF_3_^−^ or AlF_4_^−^; ref. (Bublitz et al., 2010)) (Fig. S1F; see below). Although some samples contained more than one conformational state due to incomplete conversion, distinct structures could be efficiently separated by three dimensional (3D) classifications during cryo-EM single-particle analysis.

Cryo-EM maps of ATP13A2 were reconstructed at overall resolutions of 2.5 to 3.7 Å (Fig. S1F and Tables 1 and S1). Most side chains were clearly resolved in the cryo-EM maps, enabling accurate de novo structural modeling (Fig. S2A). The reaction states of the enzyme could be assigned based on visualized densities of bound ligands and through structural comparisons with other known P-type ATPase structures (Bublitz et al., 2010; Dyla et al., 2019; Hiraizumi et al., 2019; McKenna et al., 2020; Nakanishi et al., 2020) (Fig. S2 B–M). The highest-resolution (2.5 Å) structure was obtained with WT ATP13A2 treated with AlF_4_^−^, which represents the E2-Pi state of the enzyme (Fig. S3B). However, to describe the overall architecture, we will first reference the 2.8-Å-resolution BeF_3_^−^-bound structure (Fig. 1 and S3A), because the flexible cytosolic domains are better resolved in this map. The BeF_3_^−^-bound form represents the E2P state, where the substrate-binding pocket within TMD faces toward the endo-/lysosomal interior, prior to binding a polyamine substrate.

**Table 1.** Cryo-EM data collection, refinement and validation statistics.

|  | WT<br>E2P, BeF3-bound<br>(map 3)<br>(EMD-xxxx,<br>PDB xxxx) | WT<br>E2-Pi, AIF4-bound<br>(map 6)<br>(EMD-xxxx,<br>PDB xxxx) | WT<br>E2-Pi, post-hydrolysis<br>(map 12)<br>(EMD-xxxx,<br>PDB xxxx) | WT<br>E1P-ADP<br>(map 7)<br>(EMD-xxxx,<br>PDB xxxx) | D508N<br>E1-ATP<br>(map 8)<br>(EMD-xxxx,<br>PDB xxxx) | D458N/D962N<br>E1 (apo)<br>(map 9)<br>(EMD-xxxx,<br>PDB xxxx) | D458N/D962N<br>E1 (apo)<br>(map 10)<br>(EMD-xxxx,<br>PDB xxxx) | D458N/D962N<br>E1P, AIF4-bound<br>(map11)<br>(EMD-xxxx,<br>PDB xxxx) |
| --- | --- | --- | --- | --- | --- | --- | --- | --- |
| **Data collection and processing** |  |  |  |  |  |  |  |  |
| Magnification | 64,000x | 64,000x | 64,000x | 64,000x | 64,000x | 64,000x | 64,000x | 36,000x |
| Voltage (kV) | 300 | 300 | 300 | 300 | 300 | 300 | 300 | 200 |
| Electron exposure<br>(e<sup>-</sup>/Å<sup>2</sup>) | 50 | 50 | 50 | 50 | 50 | 50 | 50 | 50 |
| Defocus range (μm) | -0.7 to -2.5 | -0.7 to -2.5 | -0.7 to -2.5 | -0.7 to -2.5 | -0.9 to -2.4 | -0.8 to -2.5 | -0.8 to -2.5 | -0.7 to -1.8 |
| Pixel size (Å) | 0.911 | 1.049 | 0.911 | 1.049 | 0.911 | 1.049 | 1.049 | 1.115 |
| Symmetry imposed | C1 | C1 | C1 | C1 | C1 | C1 | C1 | C1 |
| Initial particle images<br>(no.) | 1,437,395 | 1,311,023 | n/a | 1,311,023 | 1,431,882 | 1,181,866 | 1,181,866 | 1,406,963 |
| Final particle images<br>(no.) | 386,118 | 462,490 | 591,903 | 163,433 | 459,541 | 186,331 | 188,879 | 377,365 |
| Map resolution (Å) | 2.8 | 2.5 | 3.0 | 2.9 | 2.8 | 2.9 | 2.9 | 3.2 |
| FSC threshold | 0.143 | 0.143 | 0.143 | 0.143 | 0.143 | 0.143 | 0.143 | 0.143 |
| Map resolution<br>range (Å) | 2.5–12 | 2.4–8 | 2.6–8 | 2.5–11 | 2.4–8 | 2.4–8 | 2.5–8 | 2.8–8 |
| **Refinement** |  |  |  |  |  |  |  |  |
| Initial model used | De novo | Model 3 | Model 6 | Model 3 | Model 7 | Model 7 | Model 7 | Model 7 |
| Model resolution (Å) | 3.0 | 2.6 | 3.2 | 3.0 | 2.9 | 3.0 | 3.0 | 3.4 |
| FSC threshold | 0.5 | 0.5 | 0.5 | 0.5 | 0.5 | 0.5 | 0.5 | 0.5 |
| Map sharpening *B*<br>factor (Å<sup>2</sup>) | 75 | 63 | 116 | 56 | 99 | 79 | 77 | 117 |
| Model composition |  |  |  |  |  |  |  |  |
| Non-hydrogen<br>atoms | 7,961 | 8,113 | 8,113 | 8,026 | 8,024 | 7,887 | 7,887 | 8,006 |
| Protein residues | 1,036 | 1,034 | 1,034 | 1,040 | 1,040 | 1,040 | 1,040 | 1,043 |
| Ligands | 4 | 11 | 11 | 6 | 4 | 0 | 0 | 4 |
| *B* factors (Å<sup>2</sup>) |  |  |  |  |  |  |  |  |
| Protein | 53 | 66 | 90 | 51 | 31 | 71 | 42 | 84 |
| Ligand | 54 | 46 | 47 | 39 | 16 | - | - | 42 |
| R.m.s. deviations |  |  |  |  |  |  |  |  |
| Bond lengths (Å) | 0.004 | 0.006 | 0.009 | 0.005 | 0.005 | 0.005 | 0.004 | 0.005 |
| Bond angles (°) | 0.600 | 0.752 | 0.857 | 0.664 | 0.792 | 0.665 | 0.608 | 0.722 |
| **Validation** |  |  |  |  |  |  |  |  |
| MolProbity score | 1.43 | 1.41 | 1.61 | 1.59 | 1.53 | 1.63 | 1.55 | 1.81 |
| Clashscore | 7.67 | 7.45 | 8.84 | 8.18 | 8.18 | 8.21 | 6.02 | 10.23 |
| Poor rotamers (%) | 0 | 0 | 0 | 0 | 0 | 0.12 | 0.12 | 0.23 |
| Ramachandran plot |  |  |  |  |  |  |  |  |
| Favored (%) | 97.95 | 98.34 | 97.26 | 97.18 | 97.57 | 96.89 | 96.60 | 95.93 |
| Allowed (%) | 2.05 | 1.66 | 2.74 | 2.82 | 2.43 | 3.11 | 3.40 | 4.07 |
| Disallowed (%) | 0 | 0 | 0 | 0 | 0 | 0 | 0 | 0 |

**Figure 1.**
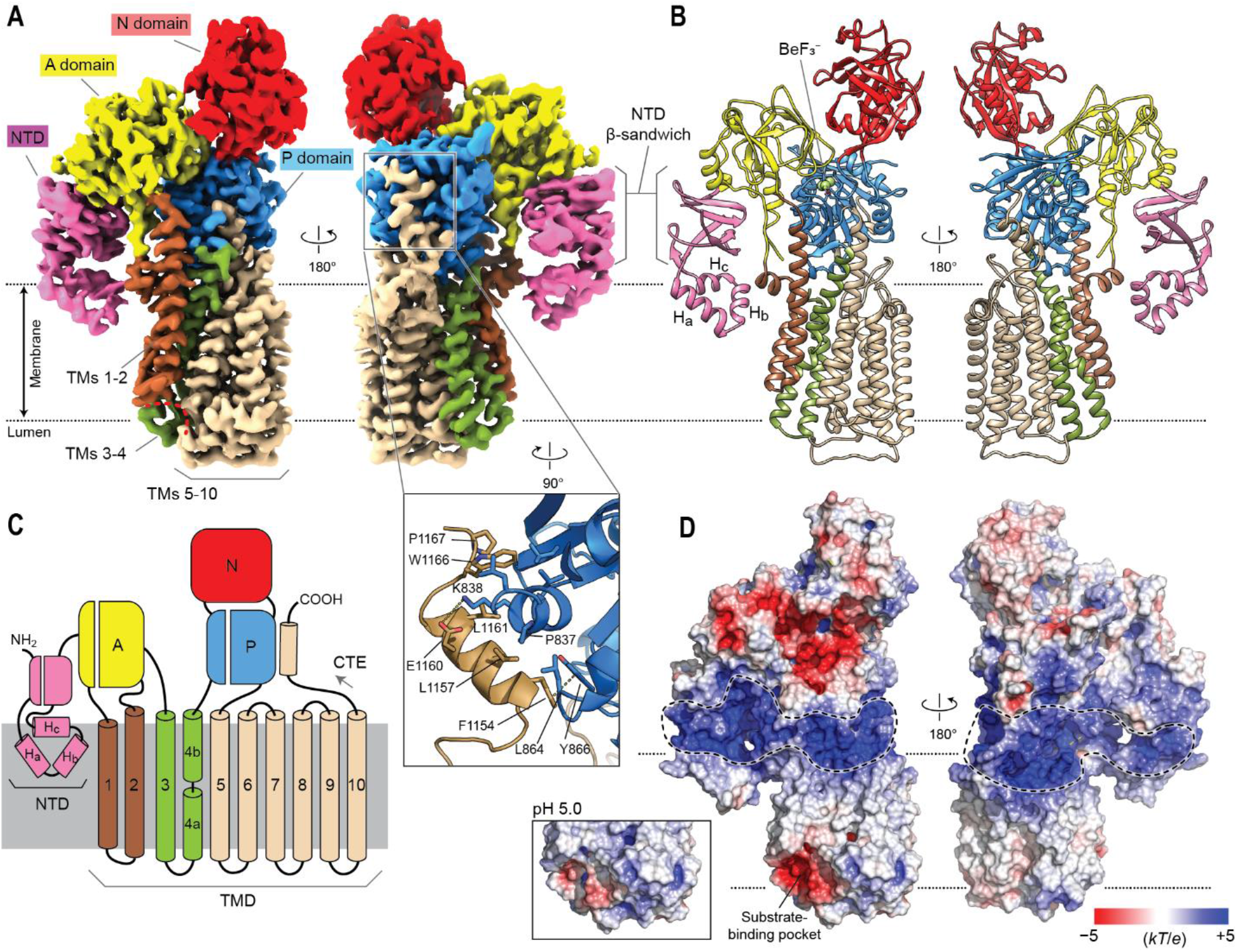
Cryo-EM structure of human ATP13A2. (**A**) 2.8-Å-resolution cryo-EM map of human ATP13A2 in the E2P-like state (BeF_3_^−^-bound form; map 3). Left, a front view; right, a view from the back. Dotted lines indicate the approximate boundaries of the endo-/lysosomal membrane. The outward-facing substrate binding pocket is indicated by a red dashed line. In the right panel, the C-terminal region is outlined by a rectangle, and its atomic model is shown in the inset. Amino acid side chains at the interface between the P domain and C-terminal extension (CTE) are shown as sticks. Dashed lines indicate interactions of K838-E1160 and Y866-Y1154. (**B**) Atomic model built into the map in A. Three membrane-embedded helices in the NTD are indicated by H_a_, H_b_, and H_c_. (**C**) Domain architecture of ATP13A2. Note that the TM4 helix is unraveled in the middle (between TM4a and TM4b) by a PP(A/V)xP motif conserved among P5B-ATPases. (**D**) Surface electrostatic potential at pH 7 was estimated and shown as a heat map. Dashed lines indicate positively charged surfaces that may interact with negatively charged lipids. Inset, electrostatics around the substrate binding pocket estimated at pH 5.

### Architecture of ATP13A2

Overall, the structure of ATP13A2 follows the canonical architecture of P-type ATPases (Fig. 1 A–C). The cytosolic domains include the actuator (A), nucleotide-binding (N), and phosphorylation (P) domains, which are responsible for converting ATP energy to conformational movements required for substrate translocation. In the membrane, ATP13A2 contains a total of ten transmembrane segments (TMs 1–10), as is common for most eukaryotic P-type ATPases (Palmgren and Nissen, 2011) (Fig. 1C).

The TMD of ATP13A2 can be divided into three groups: TMs 1–2, 3–4, and 5–10 (Fig. 1C). The substrate-binding site is formed at the interface of these three TM groups (Fig. 1A). In the BeF_3_^−^-bound E2P-like structure, the pocket is shallow and faces the interior of the endo-/lysosome (Fig. 1 A and B). Surface electrostatics calculations estimate that the pocket exhibits a negative electrostatic potential mainly due to multiple surface-exposed aspartic and glutamic acid residues (D249, E451, and D955) (Fig. 1D). While these acidic residues might be substantially neutralized in the low pH environment of the endo-/lysosomal interior, the pocket is still estimated to maintain an overall negative electrostatic potential at pH 5 (Fig. 1D, inset). These negative charges may attract positively charged polyamines into the pocket during initial polyamine binding steps.

ATP13A2 also shows unique features in both the N- and C-terminal regions (Fig. 1 A–C). An ∼180-residue-long N-terminal domain (NTD) contains three short hydrophobic α-helices (denoted H_a_, H_b_, and H_c_) that are conserved in P5B-ATPases. These helices do not traverse the membrane but instead are embedded in the cytosolic leaflet of the lipid bilayer and arranged in a triangular spade shape (Fig. 1 A– C), which is attached to the A domain by a β-sandwich. By providing an additional anchor to the membrane, this feature may play a role in guiding the conformational motions between the A domain and TMs 1–2 for substrate translocation. It may also be involved in endo-/lysosomal targeting of ATP13A2, as suggested previously (Holemans et al., 2015). In the C-terminal region, a cytosolic segment after TM10 (referred to as the C-terminal extension or CTE) forms a small helix, which is bound to the P domain mainly by hydrophobic interactions (Fig. 1A, inset). Sequence comparisons suggest that this feature is also conserved in most P5B-ATPases (Fig. S4A).

### Disease-associated mutations

The structure of ATP13A2 allowed us to map known disease-associated missense mutations in order to better understand the mechanisms of these mutations (Fig. 2A and Table S2). Many mutations were mapped in buried regions of domains, suggesting that they likely cause protein folding defects. One mutation (G517V) was mapped in the connection between the P and N domains, which may impair the interdomain movements for the catalytic reaction. The A224V and R975H mutations may affect the transport function of the enzyme by altering interactions with lipids. Interestingly, two mutations were expected to affect the functions of the NTD and CTE based on our new structures: the F177L mutation is located in the β-sandwich of the NTD (Fig. S4B), and the P836L mutation is at the interface between the P domain and CTE (Fig. 1A, inset). Similarly, a premature termination mutation (Q1135X) identified in a hereditary spastic paraplegia patent is expected to produce an enzyme lacking CTE (Estrada-Cuzcano et al., 2017).

**Figure 2.**
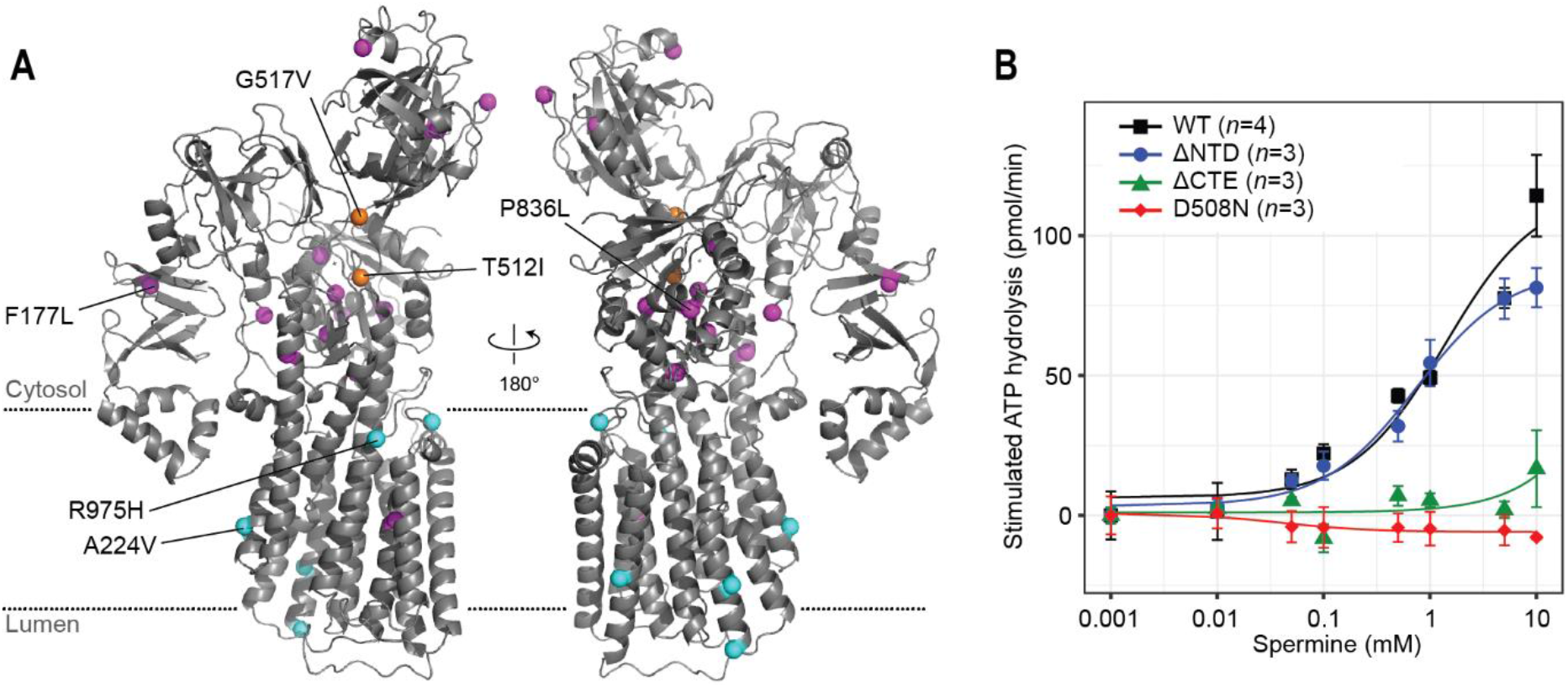
Disease-associated mutations of ATP13A2 and functional importance of CTE. (**A**) Positions of disease-associated missense mutations (colored spheres) were mapped onto the ATP13A2 E2P structure. Magenta, mutations that potentially cause protein folding defects. Orange, mutations that potentially alter the D508-phosphorylation reaction (T512I) or P-N interdomain interaction (G517V). Cyan, mutations of unknown mechanisms. (**B**) Spermine-induced ATPase stimulation of microsomes overexpressing WT ATP13A2, the D508N mutant, and mutants lacking the NTD (Δ2-179; ΔNTD) or CTE (Δ1149–1175; ΔCTE) (means and s.e.m.). Lines are fitted dose-response (Michaelis-Menten) curves.

To probe the functional importance of the NTD and CTE, we isolated microsomes overexpressing WT ATP13A2 or a mutant lacking the NTD (Δ2–179; ΔNTD) or CTE (Δ1149–1175; ΔCTE) from Sf9 cells and performed ATPase assays in the presence of varying concentrations of spermine (Fig. 2B). As previously shown (van Veen et al., 2020), the ATPase activity of WT ATP13A2 was robustly stimulated by spermine. The ΔNTD mutant showed an activity comparable to WT, suggesting that the NTD itself is not required for the catalytic cycle. Instead, the F177L mutation likely causes misfolding or decreased stability of the NTD, leading to a trafficking defect at the ER and degradation (Podhajska et al., 2012). Unlike ΔNTD, the ΔCTE mutant showed essentially no ATPase activity, despite a comparable expression level. Fluorescence size-exclusion chromatography analysis indicates a moderate tendency of the ΔCTE mutant to form high molecular-weight aggregates upon detergent extraction (Fig. S4C). While the exact function is yet to be elucidated, the CTE thus seems to be important for the activity and structural stability of ATP13A2.

### Lipid interactions

One prominent feature of the ATP13A2 structure is a belt of highly positively charged surface around the protein at the cytosol-membrane interface (Fig. 1D). This feature was not seen in other P-type ATPases, such as the sarco-/endoplasmic reticulum Ca^2+^-ATPase (SERCA), P4-flippases, and P5A-ATPase (Fig. S5A). The positive surface charge of ATP13A2 is mainly due to a high ratio of basic to acidic amino acids in this region (Fig. S5B). Homology modeling of other P5B-ATPases also showed similar distributions of charged amino acids, suggesting that this feature is common among P5B-ATPases (Fig. S5C). The extensive positive charge may recruit negatively charged lipids, including phosphatidic acid (PA) and phosphatidylinositol(3,5)biphosphate [PI(3,5)P_2_], from the cytosolic leaflet to ATP13A2. PA and PI(3,5)P_2_ have been shown to substantially stimulate the ATPase activity of ATP13A2 (Holemans et al., 2015; van Veen et al., 2020).

To test interactions between the positively charged surface and lipids, we performed 10-μs coarse-grain molecular dynamics (MD) simulations with the E2P structure and a model lysosomal membrane (Fig. S6 and Movies S1 to S4). The results show that negatively charged lipids are preferentially recruited to ATP13A2, in particular phosphoinositide (PI) (Fig. S6 A and B). If present, the polyanionic PI(3,5)P_2_ lipid strongly accumulated around ATP13A2, outcompeting less negatively charged lipids (Fig. S6 C and D). A similar result was also obtained with an E2-Pi structure (Fig. S6 E–H). As hypothesized, the positive electrostatic belt thus causes strong lipid sorting in the simulations and enhances the local concentration of (poly)anionic lipids around ATP13A2.

### Comparison with the P5A-ATPase

P5A- and P5B-ATPases share substantial sequence similarity (∼25% identity), and yet they have vastly different substrate specificities. The P5A-ATPase extracts mis-inserted transmembrane proteins by flipping an ER-lumenal hydrophilic polypeptide segment (length of up to 7–10 residues; ∼1,000 Da) attached to a transmembrane helix into the cytosol (McKenna et al., 2020). By contrast, polyamines are water-soluble and considerably narrower in diameter (molecular weight of spermine is ∼200 Da). Accordingly, the structures of the substrate-binding pockets of the two ATPases were expected to be quite different.

A structural comparison between ATP13A2 and *Saccharomyces cerevisiae* P5A-ATPase Spf1 (a homolog of the mammalian ATP13A1) (McKenna et al., 2020) explains how a relatively small structural difference dramatically changes the size and topology of the substrate-binding pocket (Fig. 3). As expected from their sequence similarity, the overall structures of ATP13A2 and Spf1 are similar, including in the arrangements of their TMs (Fig. 3 A–C). However, a major difference lies in the position of TMs 5–10, which are rotated away from the other TMs by ∼20° along TM5 in Spf1 compared with ATP13A2 (Fig. 3 C–F). This rotation causes a widening of the substrate-binding pocket in Spf1 and creates a laterally open topology to accommodate and flip a polypeptide segment attached to a lipid-embedded TM helix. By contrast, TMs 5–10 of ATP13A2 are positioned adjacent to TMs 1–2, forming a narrow cavity that is separated from the lipid phase and suited for a small, charged molecule.

**Figure 3.**
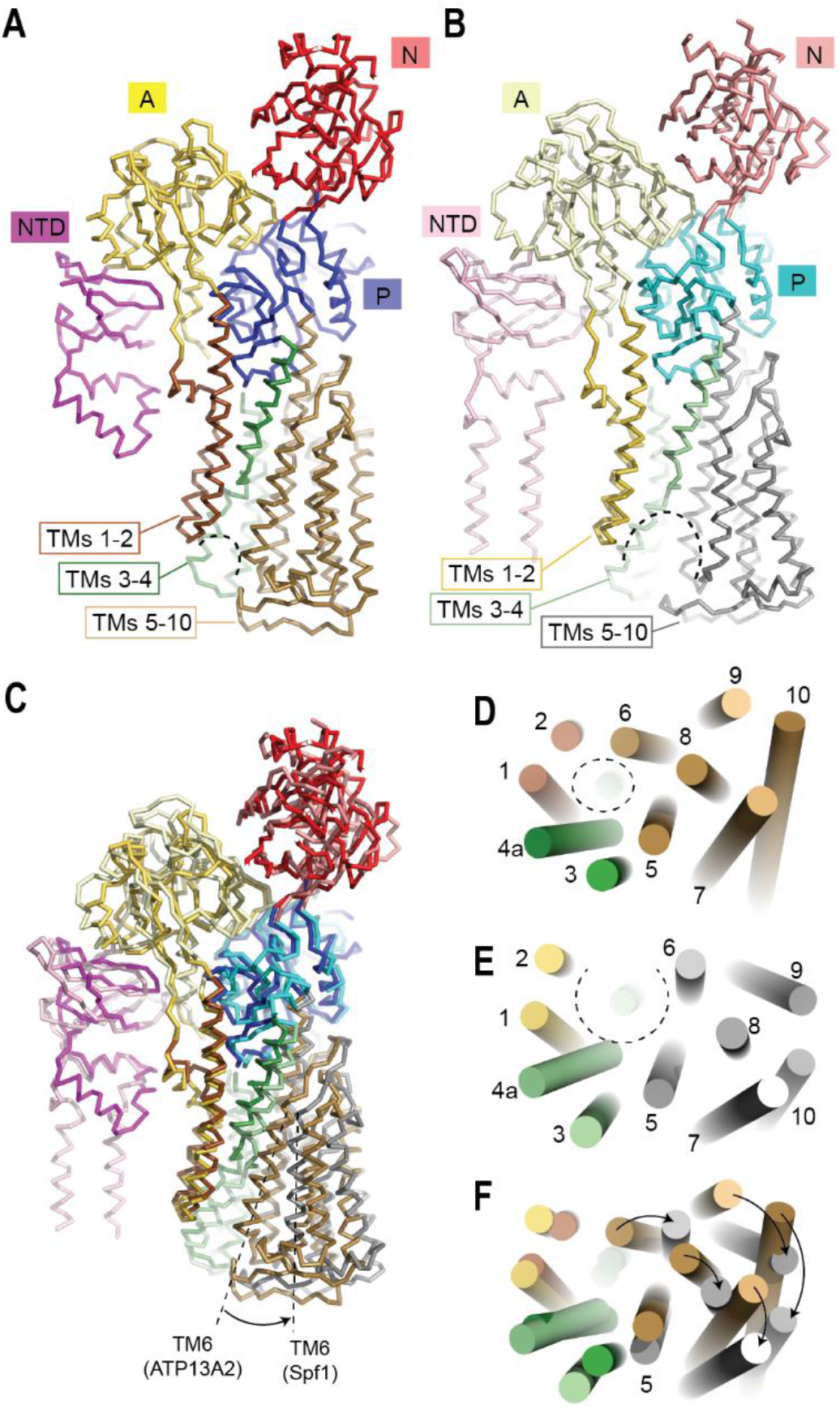
Structural comparison between ATP13A2 and the P5A-ATPase Spf1. (**A**) Structure of human ATP13A2 in the E2P-like state shown in a backbone-trace representation. The color scheme is the same as Fig. 1 A–C. The substrate binding pocket is indicated by a dashed line. (**B**) As in A, but with a BeF_3_^−^-bound E2P-like structure of Spf1 from *S. cerevisiae*. (**C**) The structures in A and B were aligned based on the P domain and TMs 1–4. Positions of TM6 are indicated. (**D**) TMD arrangement of ATP13A2 shown in a view from the lumen. The substrate-binding pocket is indicated by a dashed circle. (**E**) As in D, but with the Spf1 structure. Note that the substrate-binding pocket of Spf1 is open toward both the ER lumen and lipid phase of the membrane. (**F**) Overlay between D and E. Rotational rearrangement of TMs 5-10 along TM5 is indicated by arrows.

### Structural basis of polyamine binding

To understand the molecular basis of ATP13A2’s polyamine specificity, a polyamine-bound ATP13A2 structure needed to be determined. Serendipitously, a strong, well-resolved polyamine-like density could be seen in the substrate-binding pocket of the E2-Pi structure without supplementing exogenous polyamines (Fig. 4 A–C and S3B). The polyamine-bound E2-Pi conformation was the predominant form of the WT enzyme in our protein preparations, as it was observed in all WT ATP13A2 samples.

**Figure 4.**
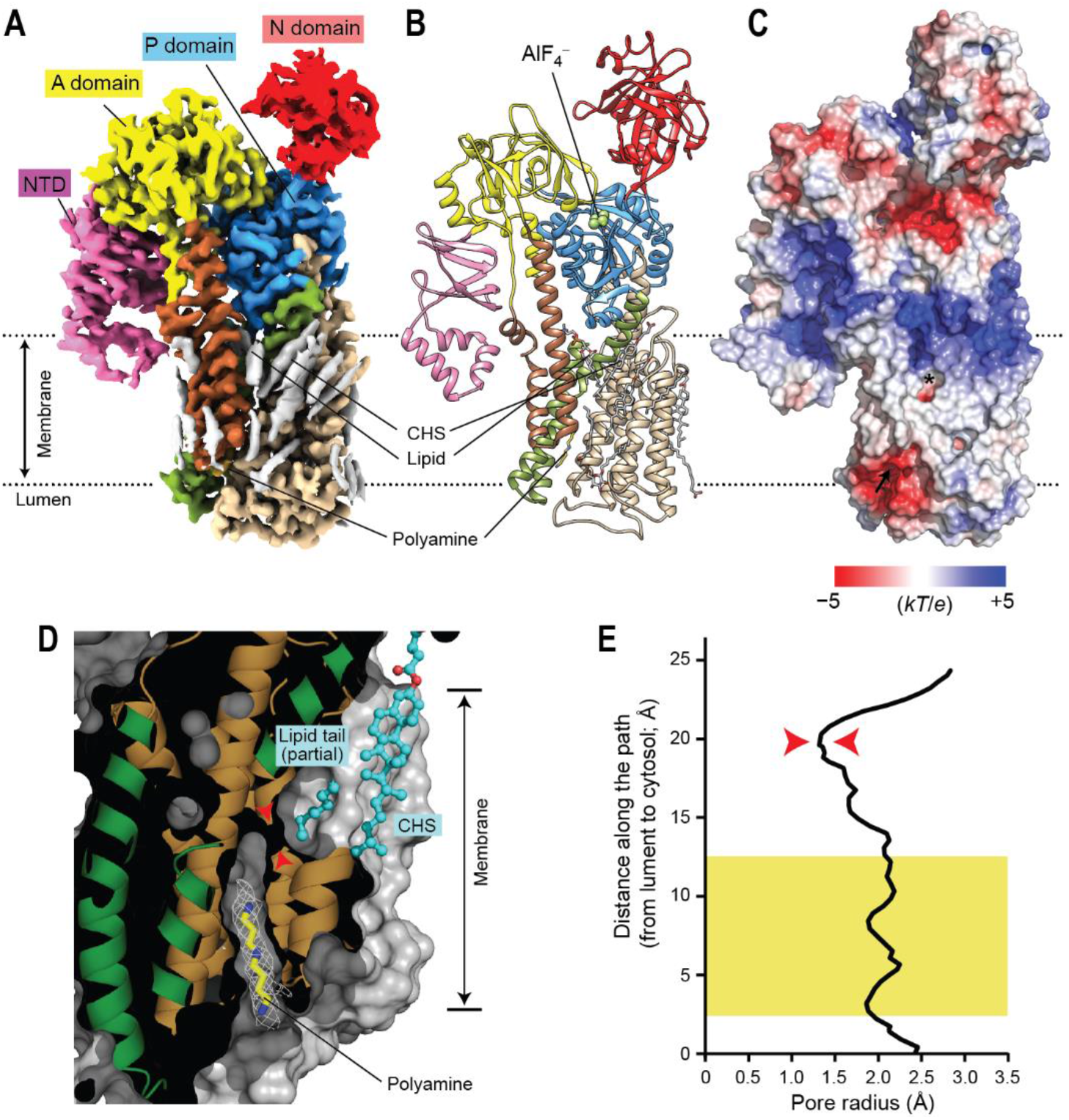
Polyamine-bound E2-Pi structure of ATP13A2. (**A** and **B**) 2.5-Å-resolution cryo-EM map (A) and atomic model (B) of ATP13A2 in the polyamine-bound E2-Pi-like state (AlF_4_^−^-bound form; map 6). The domains are colored as in Fig. 1A. Gray densities in A are lipids and detergent molecules. (**C**) Surface electrostatic potential calculated at pH 7. Arrow, polyamine entrance; asterisk, putative polyamine exit (see panel D). (**D**) Cutaway view of the substrate-binding pocket in the E2-Pi structure (protein surface shown in gray). The view angle is an ∼90° right-hand-rotated with respect to that of panels A–C. The observed polyamine density is shown as a white mesh. The lipid tail and CHS molecule that sits in front of the putative polyamine exit are shown as cyan balls and sticks. Red arrowheads indicate the narrowest neck of the cavity. Approximate membrane boundaries are also indicated. (**E**) Radius profile of the transport conduit. The region marked by a yellow box is the position of the polyamine-binding site. Red arrowheads indicate the narrowest (1.3 Å in radius) region corresponding to the region marked in the right panel of D.

Particularly, when purified in the presence of AlF_4_^−^ or without any added factors, WT enzyme was found to be almost exclusively in the E2-Pi conformation (Fig. S1F). More specifically, the AlF_4_^−^-bound WT enzyme is likely in the dephosphorylation transition state, whereas the other observed E2-Pi structures may represent the product (post-hydrolysis) state (Bublitz et al., 2010). However, the conformations in these two states seem indistinguishable (Fig. S2 B–E, G, and H), similar to SERCA (Bublitz et al., 2010; Dyla et al., 2019). In the E2-Pi state, ATP13A2 seems to tightly bind polyamines, which in our samples must have originated from insect cells or the culture medium.

Compared with the E2P structure, the E2-Pi structure shows a substantially deeper substrate-binding cavity that is well-suited to bind an elongated polyamine molecule (Fig. 4D). This channel-like cavity forms a continuous conduit through the TMD but is constricted to ∼1.3 Å in radius near the cytosolic end, which would be too narrow for the polyamine molecule to readily pass (Fig. 4D). Thus the conduit is blocked on the cytosol side. However, the topology of the detected conduit suggests that the polyamine exit may be formed within the hydrophobic layer of the cytosolic leaflet of the membrane (Fig. 4 C and D). In our E2-Pi structures, this opening is further capped by a lipid acyl chain and a CHS molecule (Fig.4D).

The bound polyamine molecule, which we assigned as spermidine based on the length of the density feature, is in an extended conformation and positioned along the cavity in the endo-/lysosomal lumenal leaflet. The cavity is sufficiently long (∼15 Å) to accommodate the tetraamine spermine but is only wide enough (∼2 Å in radius) to accommodate one extended polyamine molecule (Fig. 4E). Thus, when the cavity is occupied by a polyamine, other molecules such as water or ions would not be able to pass.

Binding of the polyamine is mediated largely by a network of ionic and cation-π interactions between all three amine nitrogen atoms of the spermidine molecule and the acidic and aromatic side chains that line the cavity (Fig. 5 A and B). Specifically, the middle amine nitrogen atom of spermidine is coordinated by Y251, D458, and F958, and the two terminal amines are coordinated mainly by D249 and D955 on one side and D962 on the other. These amino acids are strongly conserved among P5B-ATPases, suggesting that the same or a similar mode of interaction may be used for substrate binding in other P5B-ATPases. This arrangement of acidic and aromatic amino acids and their interactions with the polyamine substrate in ATP13A2 are strikingly analogous to those observed in bacterial periplasmic polyamine-binding proteins, although their protein folds are completely different (Kashiwagi et al., 1996; Sugiyama et al., 1996) (Fig. 5 C and D). This suggests convergent evolution in polyamine recognition mechanisms.

**Figure 5.**
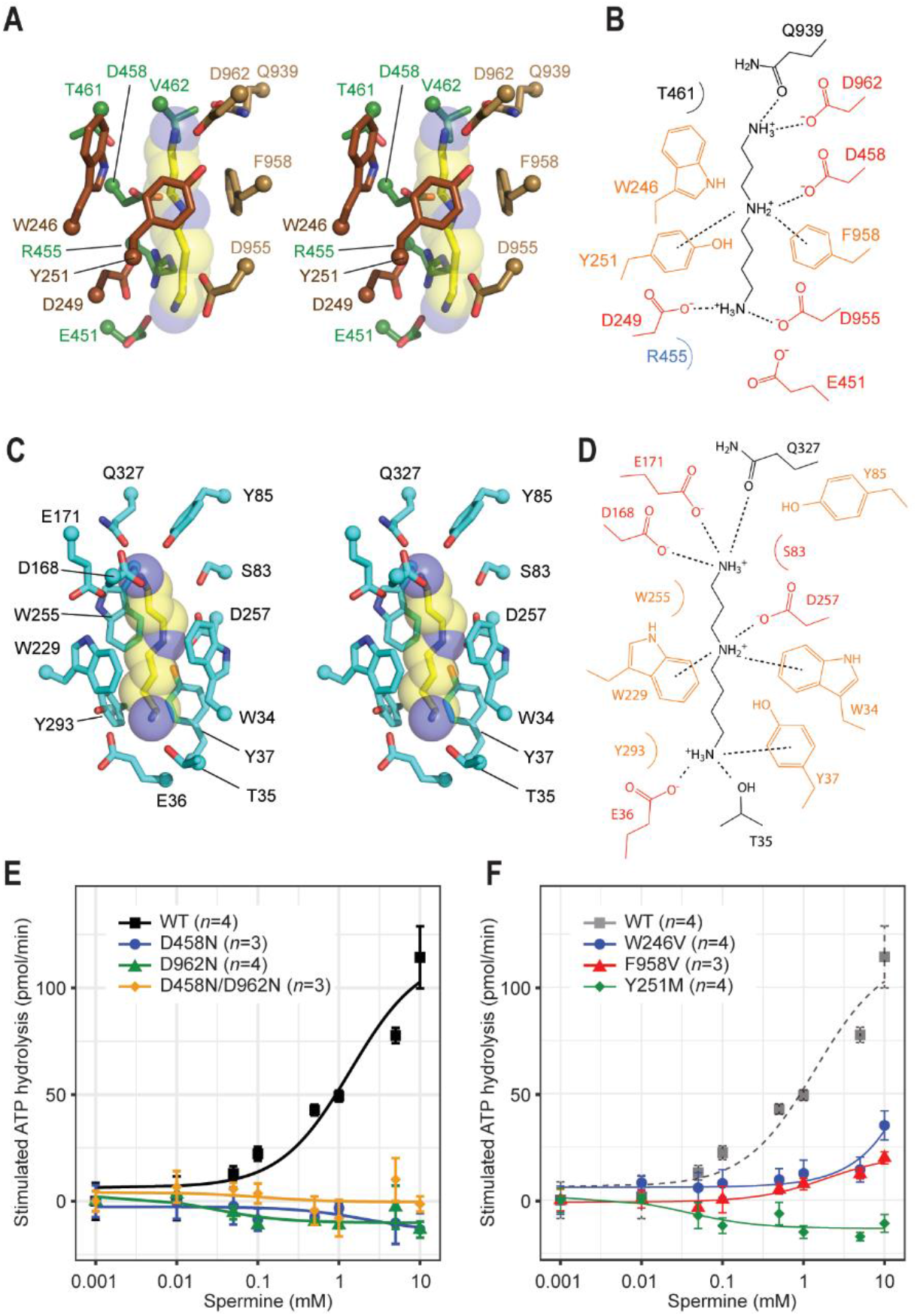
Structural basis of polyamine binding in ATP13A2. (**A**) Stereo view of the polyamine-binding site of ATP13A2 in the E2-Pi structure. The modelled spermidine molecule is shown as yellow (carbon)/blue (nitrogen) sticks and semitransparent spheres. Side chains of ATP13A2 coordinating the spermidine molecule are shown in a stick representation (brown, TMs 1–2; green TM4; tan, TM6). (**B**) Diagram showing interactions between ATP13A2 and the polyamine (spermidine) molecule. Ionic and cation-π interactions are depicted as dashed lines. (**C**) As in A, but with the polyamine-binding site of *Escherichia coli* PotD (PDB: 1POT). (**D**) As in B, but for the PotD-spermidine structure. (**E**) Spermine-induced ATPase stimulation of microsomes expressing WT ATP13A2 or mutants of polyamine-interacting aspartate residues (means and s.e.m.). Lines are fitted dose-response curves. (**F**) As in E, but with mutants of polyamine-interacting aromatic residues.

We tested the effect of mutating these acidic and aromatic amino acids on the ATPase activity of ATP13A2 in the presence of spermine (Fig. 5 E and F). As expected, the D458N, D962N, and D458N/D962N mutants showed no spermine-dependent ATPase stimulation, suggesting their inability to bind spermine. Similarly, the W246V, Y251M, and F958V mutants also displayed little or no ATPase stimulation. These results confirm the importance of these amino acids for polyamine binding.

### Conformational changes in TMD and putative polyamine paths

Next, to understand the mechanism of polyamine transport by ATP13A2, we sought to determine structures of its E1 states. Many P-type ATPases, including the P5A-ATPase, display an inward-open (cytosolically open) conformation in the E1 state (Bai et al., 2020; McKenna et al., 2020; Palmgren and Nissen, 2011; Toyoshima et al., 2000; Winther et al., 2013). Typically, in other P-type ATPases, the E1-apo state can be obtained simply by preparing the WT enzyme in the absence of a nucleotide or phosphate analog (Bai et al., 2020; Hiraizumi et al., 2019; McKenna et al., 2020; Nakanishi et al., 2020). However, WT ATP13A2 purified in this way was found predominantly in a polyamine-bound E2-Pi conformation (Fig. S1F). Addition of nonhydrolyzable ATP analog AMP-PCP to this form did not change the state, likely because AMP-PCP was unable to bind (Figs. S1F and S2C). In cells, WT ATP13A2 probably assumes an E1P or E2P state (Holemans et al., 2015), and tight binding of polyamines during detergent extraction at a neutral pH might have converted ATP13A2 into the E2-Pi state. To prevent this conversion and shift the equilibrium to favor an E1P-ADP state, we treated WT ATP13A2-expressed membranes with ADP prior to extraction. Indeed, this method yielded an E1P-ADP-like structure from ∼20% particles (Fig. S1F). In separate approaches to obtain additional E1 states, we also analyzed mutants defective in phosphorylation (D508N) or polyamine binding (D458N/D962N). The D508N mutant in the presence of ATP exclusively yielded an E1-ATP-like structure, whereas the D458N/D962N mutant yielded an E1 (apo) structure in the absence of a ligand and an E1P-like structure in the presence of AlF_4−_ (Fig. S1F). A lack of an E2-Pi population in the D458N/D962N mutant, in stark contrast to WT ATP13A2, suggests that polyamine binding is necessary to form the E2-Pi state. Thus, this finding explains the dramatic stimulation of ATP13A2’s ATPase activity induced by polyamines (van Veen et al., 2020).

All E1 structures of ATP13A2 display a highly similar overall conformation, including that of the TMD (Fig. S7A). One exception is the flexible N domain, which either is in a closed position when ATP or ADP is bound at the interface between the N and P domains, or swings outward in the absence of adenine nucleotide (Fig. S7 A and B). The lack of major conformational changes in the TMD between the E1 structures and the nucleotide-dependent movement in the N domain are consistent with observations made previously with other P-type ATPases (Bai et al., 2020; Dyla et al., 2019; Hiraizumi et al., 2019; McKenna et al., 2020; Nakanishi et al., 2020).

Importantly, the TMD in the E1 states displays a conformation that is clearly distinct from the E2P and E2-Pi structures (Fig. 6, S7 C and E). Unlike E2-Pi, the substrate cavity in the E1 states is too narrow (with a radius of ∼1 and 1.5Å) to accommodate a polyamine (Fig. 6 A–C). In fact, the cavity seems inaccessible from both sides. Thus, the conformation of the TMD in E1 states represents a “occluded” state. Nevertheless, like the E2-Pi structure, a continuous conduit with a potential polyamine exit within the membrane could be detected in the E1 structures (Fig. 6 A and B; cyan path), suggesting that the bound polyamine molecule may follow this path to exit to the cytosol during the E2-Pi to E1 transition. Alternatively, the polyamine molecule may travel through another path branching out from the above-mentioned conduit around Y254 (Fig. 6 A and B; gray path). In our structures, this path is blocked by the side chains of Q239, I258, V464, and P466, but we cannot rule out the possibility that it may open during the E2-Pi to E1 transition.

**Figure 6.**
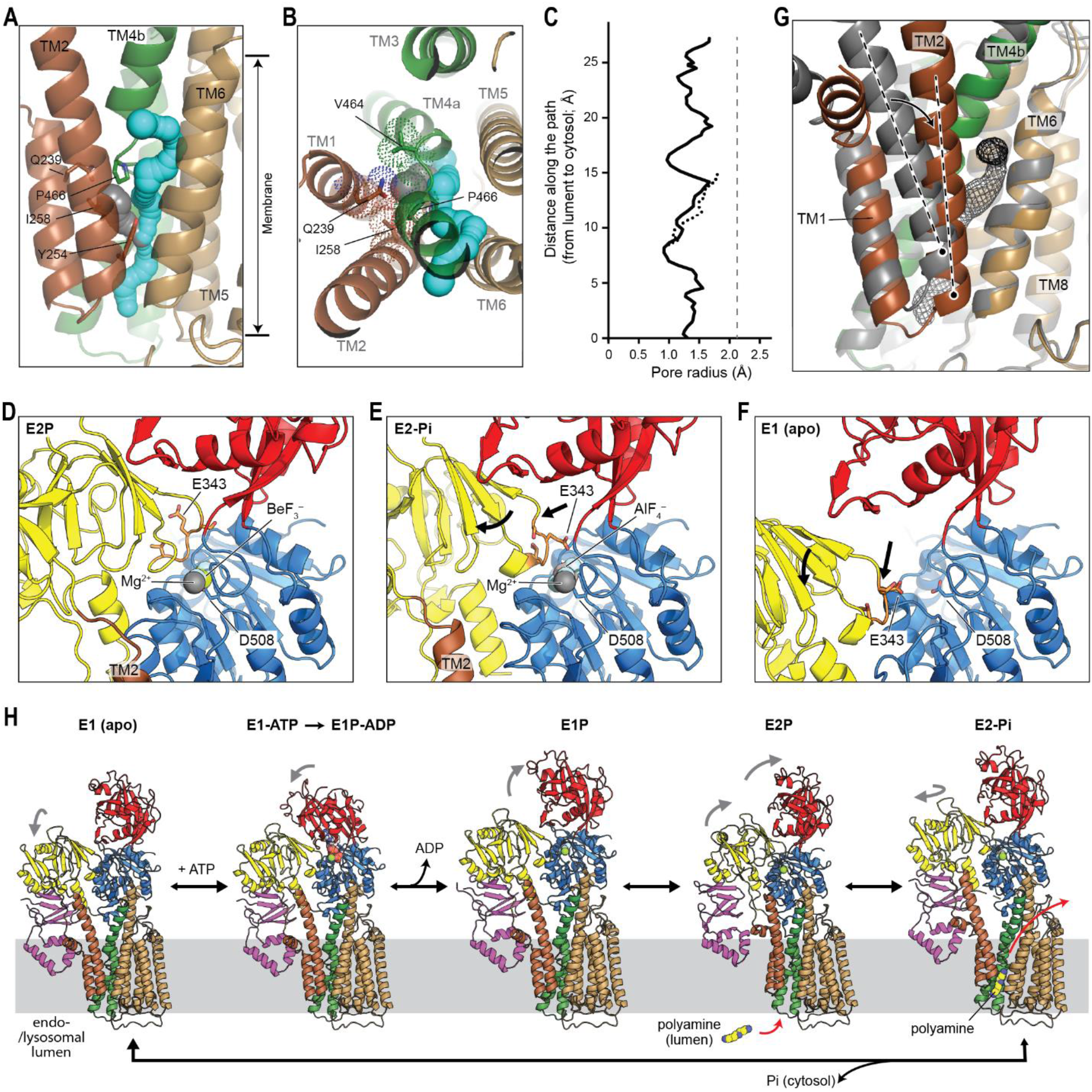
Structure of ATP13A2 in the E1 state and conformational transitions. (**A** and **B**) Structure of the TMD of D458N/D962N ATP13A2 in the E1 (apo) state. Shown are front (A) and top (B) views. TMs are colored in the same scheme as Figs. 1 and 2. The potential polyamine paths were detected by a spherical probe and are shown in cyan and gray. Approximate membrane boundaries are indicated. The side chains of the indicated amino acids are shown in sticks and dots (panel B only). (**C**) Radius profiles of the paths shown in A and B. The solid and dotted lines correspond to the cyan and gray paths, respectively. The gray dashed line indicates an average pore radius (2.1 Å) of the polyamine-bound region in the E2-Pi structure (see Fig. 4E). (**D**) View into the phosphorylation site (D508) in the E2P structure. A, P, and N domains are colored yellow, blue, and red, respectively. The TGES loop containing E343 is in orange with side chains shown in a stick representation. (**E**) As in D, but with the E2-Pi structure. Relative overall movements of the TGES loop and A domain from the positions in the E2P state are indicated by black arrows. (**F**) As in D and E, but with the E1 (apo) structure. Relative overall movements of the TGES loop and A domain from the positions in the E2-Pi state are indicated by black arrows. (**G**) Conformational transition of TMs 1–2 (indicated by an arrow) from the E2-Pi (in gray) to E1 (apo) state (in color). The structures were aligned with respect to TMs 5–10. The axis of TM2 is indicated by a dashed line. The putative polyamine transport conduit (cyan path in panels A and B) is shown in gray mesh. (**H**) Polyamine-transport cycle of ATP13A2. All displayed structures were experimentally observed in this study. Gray arrows indicate major movements of the cytosolic domains (with respect to the previous state). Also see Fig. S7.

Like other P-type ATPases, the transitions between the different TMD conformations in ATP13A2 are mediated largely by movements of the A domain with respect to the P domain, which are relayed into conformational changes between TMs 1–2 and the rest of the TMD (Fig. 6 E–G). The A domain contains the conserved Thr^341^-Gly-Glu-Ser^344^ (TGES) loop, where the side-chain carboxyl group of E343 catalyzes dephosphorylation. In both E2P and E2-Pi states, the TGES loop is in close proximity to the phosphorylated D508 of the P domain. However, during the E2P to E2-Pi transition, the A domain is tilted by ∼20°, because the TGES loop is displaced by ∼4 Å as the side chain of E343 becomes directed toward the aspartyl phosphate for dephosphorylation (Fig. 6 D and E). This motion converts TMD from the outward-facing (E2P) to the channel-like (E2-Pi) conformation (Fig. S7D). In the E1 structures, the TGES loop is further displaced (∼13 Å) from D508 in an orthogonal direction (Fig. 6F). This leads to a translational (∼5 Å downward) and rotational (∼12° towards the front) movement of TMs 1–2 (Figs. 6G and S7E), causing a narrowing of the substrate conduit in the E2-Pi to E1 transition.

## Discussion

Our study demonstrates that ATP13A2 is a dedicated polyamine transporter and reveals the molecular basis for its polyamine recognition and transport. Our structural analysis captured nearly all major intermediate states of the ATP13A2 transport cycle. This allows us to propose a tentative working model for polyamine export by ATP13A2 on the basis of the Post-Albers scheme (Fig. 6H). In this model, polyamine binding from the end-/lysosomal lumen and membrane translocation occurs during the transitions from the E2P to E2-Pi state and from the E2-Pi to E1 state, respectively. In the E2-Pi state, the TMD of ATP13A2 forms an elongated conduit of ∼2-Å radius, which binds a polyamine molecule in an extended conformation with high affinity (Fig. 4D). During the transition from the E2-Pi to E1 state, the polyamine molecule would be released to the cytosol. The conformational trajectory of this step remains unclear as our current analysis did not resolve further intermediates between the E2-Pi to E1 transition (e.g., E2). While this transition would likely involve a transient luminally-closed but cytosolically-open state, the formation of a large cytosolic opening seems unlikely, considering the closed conformation observed in the E1 states.

The narrow, channel-like conduit and the mode of polyamine binding seen in the E2-Pi structure suggest that ATP13A2 may deviate from the canonical model of active transport. Instead of simply transitioning between outward-open, occluded, and inward-open conformations, polyamine transport by ATP13A2 likely requires the polyamine molecule to move within the narrow and elongated cavity from the lumenal entrance toward the cytosolic exit. Our structures suggest that without such movement, the lumenal gate would not be able to close, being blocked by the polyamine molecule near the cavity entrance. The motion of polyamine away from the lumen might be driven partly by the endo-/lysosomal membrane potential, in addition to a local electrostatic gradient within the cavity and conformational changes of the cavity during the E2-Pi to E1 transition. Similar voltage-dependent passage of polyamines along pores of ion channels has been well documented (Twomey et al., 2018; Williams, 1997). A gradual upward movement would also explain how ATP13A2 binds and transports polyamines of different lengths.

The exact path through which the bound polyamine molecule travels to the cytosol remains to be elucidated. However, our structures raise a possibility that the polyamine exit may be formed within the cytosolic leaflet of the membrane. In this scenario, the polyamine molecule would move along the protein-lipid interface toward the cytosol as it exits through the opening (Fig. S7F). The membrane might locally thin around the exit site, which would facilitate the solvation of polyamine charges. The exiting polyamine molecule would also likely interact with (poly)anionic lipids (Yung and Green, 1986) recruited to the surface of ATP13A2 (Fig. S6), which may promote the release of the polyamine molecule. This mechanism could enhance efficiency and directionality of polyamine export because once completely released into the cytosol, the polyamine would be blocked from returning to the cavity by a seal of lipids. This valve-like mechanism would also help prevent ion leakage by minimizing their passage through the conduit, which could dissipate the endo-/lysosomal membrane potential and proton gradient.

In addition to P5B-ATPases, putative polyamine transporters have been identified in other membrane transporter families, such as the ATP-binding cassette (ABC) transporters and solute carriers (SLC) (Aouida et al., 2005; Fujita et al., 2012; Hiasa et al., 2014; Igarashi and Kashiwagi, 1999). To our knowledge, none of these transporters has been structurally characterized. While their transport mechanisms remain to be elucidated, the convergent evolution of polyamine-binding modes we identified here (Fig. 5 A–D) suggests that they may use similar mechanisms for substrate recognition.

## Acknowledgments

We thank D. Toso and J. Remis for support for electron microscope operation, and J. Hurley and R. MacKinnon for critical reading of manuscript. This work was supported by the Vallee Scholars Program (E.P.) and Pew Biomedical Scholars Program (E.P.). S.I.S. was supported by an NIH training grant (5T32GM008295) and an NSF Graduate Research Fellowship (DGE 1752814). S.v.B and G.H. thank the Max Planck Society for support.

## Author contributions

E.P. conceived and supervised the project. S.I.S. prepared cryo-EM samples and performed biochemical assays. E.P. and S.I.S. collected and analyzed cryo-EM data, built atomic models, interpreted results. S.v.B. and G.H. performed MD simulations and interpreted results. E.P. and S.I.S. wrote the manuscript with input from all authors.

## Competing interests

The authors declare no competing interests.

## Methods

### DNA constructs

A coding sequence for full-length human ATP13A2 (isoform2; NCBI sequence ID NP_001135445.1; UniProt ID Q9NQ11-3) was obtained by PCR amplification of clones (Clone IDs: 5240813 and 5139467) from Mammalian Gene Collection cDNAs (Horizon Discovery) and inserted into pFastBac-1 (Thermo Fisher Scientific) by standard molecular biology techniques. The coding sequence is followed by an HRV 3C protease site and an enhanced GFP tag. Mutant ATP13A2 constructs were generated by PCR, and sequences were verified by Sanger DNA sequencing.

### Baculovirus generation and Sf9 expression

Recombinant bacmids were generated from the pFastBac1 constructs using the Bac-to-Bac baculovirus expression system. DH10Bac *E. coli* competent cells (Thermo Fisher Scientific) were transformed with each pFastBac1-ATP13A2-GFP construct, and colonies were selected on an LB agar plate containing 50 μg/mL kanamycin, 7 μg/mL gentamycin, 10 μg/mL tetracycline, 100 μg/mL Bluo-Gal, 40 μg/mL IPTG. Bacmids were isolated and used for transfection of Sf9 cells. Sf9 cells were cultured in ESF 921 medium (Expression Systems) at 27 °C.

Baculovirus were generated by transfecting Sf9 cells using Cellfectin II reagent (Thermo Fisher Scientific), according to the standard Bac-to-Bac protocol or linear polyethylenimidine (PEI MAX; Polysciences) (Scholz and Suppmann, 2017). When necessary, baculovirus was amplified by infecting Sf9 cells at a 1:1000 volume-to-volume ratio and harvesting the culture supernatant after 4 days post-infection. For protein expression, Sf9 cells were infected at ∼1.5–2.0 x 10^6^ cells/mL. Expression of ATP13A2 was monitored by GFP fluorescence of infected cells. Cells were harvested by centrifugation (1,500*g* for 7 min), 2–3 days post-infection, before any substantial cell death. Cell pellets were frozen in liquid nitrogen and stored at −80 °C until use.

### Preparation of microsomes

Sf9 cell pellets expressing GFP-tagged ATP13A2 constructs from 0.4 L cultures were thawed and washed twice with 15 mL of ice-cold phosphate-buffered saline (pH 7.4) (137 mM NaCl, 2.7 mM KCl, 10 mM Na_2_HPO_4_, 1.8 mM KH_2_PO_4_). All subsequent steps were carried out at 4 °C. Cells were collected by centrifugation (1,500*g* for 7 min) and resuspended in 10 mL of hypotonic lysis buffer (10 mM Tris pH 7.5, 0.5 mM MgCl_2_, 2 mM DTT, 0.1 mM PMSF, 5 µg/mL aprotinin, 5 µg/mL leupeptin, and 1 µg/mL pepstatin A). Cells were swollen in the lysis buffer by incubating them on ice for 10–15 min and lysed with 40 strokes of a Dounce homogenizer. The lysate was diluted using 10 mL of resuspension buffer (10 mM Tris pH 7.5, 0.5 M sucrose, 0.3 M NaCl, 2 mM DTT, and 0.5 mM PMSF) and further homogenized with 20 strokes of a Dounce homogenizer. Subcellular fractionation was subsequently performed through steps of differential centrifugation. The suspension was centrifuged at 1,000*g* for 4 min to remove unbroken cells and nuclear debris, and the supernatant was centrifuged at 10,000*g* for 20 min (Sorvall SS-34 rotor). The supernatant from this step was then subjected to ultracentrifugation at 200,000*g* for 35 min (Beckman Type 45 Ti rotor) to collect microsomes. The microsomal pellet was resuspended in 0.25 M sucrose containing 5 µg/mL aprotinin, 5 µg/mL leupeptin, 1 µg/mL pepstatin A, and 2 mM PMSF at a concentration between 2 and 5 mg/mL. Microsome concentrations were measured based on the total protein concentration using the Bradford assay with bovine serum albumin as a standard. Aliquots of the resuspended microsomes were frozen in liquid nitrogen and stored at −80 °C until use.

### Fluorescence size exclusion chromatography (FSEC)

100 µg microsomes were solubilized in a lysis buffer containing 50 mM Tris pH 7.5, 200 mM NaCl, 1 mM EDTA, 1 mM DTT, 10% glycerol, 5 µg/mL aprotinin, 5 µg/mL leupeptin, 1 µg/mL pepstatin A, and 2 mM PMSF, 1% n-dodecyl-β-D-maltopyranoside (DDM; Anatrace), and 0.2% cholesteryl hemisuccinate (CHS; Anatrace) for 2–3 h at 4 °C. Solubilized microsomes were clarified by centrifugation (17,000*g* for 1 h at 4 °C). The samples (equivalent to ∼85 µg microsomes) were then analyzed with an HPLC system and injected to a Superose 6 column (GE Life Sciences), equilibrated in a running buffer containing 25 mM Tris pH 7.5, 100 mM NaCl, 1 mM EDTA, 0.03% DDM, 0.006% CHS. Elution of the GFP-tagged ATP13A2 constructs was monitored by a fluorometer (λ_ex_= 475 nm; λ_em_= 510 nm; with a fixed gain) connected to the column.

### ATPase activity assay

The ATPase activity of ATP13A2 in microsomes was measured using a luminescence-based ADP detection assay (ADP Glo Max assay; Promega). Reactions were performed in a final volume of 5 µL containing 50 mM MOPS-KOH pH 7.0, 100 mM KCl, 11 mM MgCl_2_, 1 mM DTT, 0.02% DDM, 1 μg microsomes, 5 mM ATP, and various concentrations of spermine (SPM). Before initiating the reaction by addition of ATP, microsomes were first allowed to equilibrate in the reaction buffer for 1.5 h on ice and for 10 min at 37 °C. The ATPase reaction was performed for 15 min at 37 °C upon addition of ATP and was terminated by heating samples for 5 min at 80 °C. The reaction was then mixed with 5 µL of ADP-Glo Reagent for 60–80 min at 23°C, followed by addition of 10 uL of ADP-Glo Max Detection Reagent for 60 min. Luminescence was measured in a 384 multi-well plate using a luminometer (NOVOstar; BMG Labtech). Raw luminescence values were converted to ADP amounts based on a standard curve generated with 5 mM ATP/ADP mixtures at varying ratios but without microsomes. Averages and s.e.m. were calculated. Averages were corrected for the basal ATP hydrolysis activities of microsomes by subtracting the value at 0.001 mM SPM. Dose-response curves were generated using the R software with the *drc* (Ritz et al., 2015) and *ggplot2* (https://ggplot2.tidyverse.org) packages.

### Purification of human ATP13A2 for cryo-EM analysis

Sf9 cells expressing GFP-tagged ATP13A2 constructs were thawed and resuspended in a lysis buffer (Buffer L1) containing 50 mM Tris pH 7.5, 200 mM NaCl, 1 mM EDTA, 1 mM DTT, 10% glycerol, 5 µg/mL aprotinin, 5 µg/mL leupeptin, 1 µg/mL pepstatin A, and 2 mM PMSF. All subsequent steps were carried out at 4 °C. Cells were mechanically lysed by douncing and unbroken cells, and large debris was removed by centrifugation at 4,000*g* for 10 min. Membranes were pelleted by ultracentrifugation (Beckman Type 45 Ti rotor) at 100,000*g* for 1.5 h and resuspended in Buffer L1. The membranes were then solubilized with 1% DDM and 0.2% CHS for 2.5 h. The lysate was clarified by ultracentrifugation (Beckman Type 45 Ti rotor) at 100,000*g* for 1 h, and the supernatant was mixed with Sepharose beads conjugated with anti-GFP nanobody for 2.5 h. The beads were washed with approximately 30 column volumes of a wash buffer (Buffer W1) containing 25 mM Tris pH 7.5, 100 mM NaCl, 1 mM EDTA, 1 mM DTT, 0.03% DDM, 0.006% CHS. ATP13A2 was eluted by incubating the beads with ∼10 µg/mL HRV 3C protease (in Buffer W1) for ∼14 h. The eluate was concentrated using Amicon Ultra (cut-off 100k; GE Life Sciences) and injected into a Superose 6 Increase 10/300 GL column (GE Life sciences), equilibrated with Buffer W1. Peak fractions were pooled and concentrated to ∼5–7 mg/mL before cryo-EM grid preparation. To obtain ATP13A2 in specific reaction intermediate states, this base procedure was modified as described below.

The WT ATP13A2 sample that yielded map 1 was purified without isolating crude membranes. Sf9 cells were directly lysed and solubilized with Buffer L1, 1% DDM, and 0.2% CHS for 3 h. All subsequent steps were the same as the base procedure.

For WT ATP13A2 prepared with AMP-PCP (sample for map 2), the crude membrane pellet was resuspended in a lysis buffer (Buffer L2) containing 50 mM Tris pH 7.5, 200 mM NaCl, 1 mM EDTA, 1 mM DTT, 10% glycerol, 10 mM MgCl_2_, 0.5 mM spermine (SPM; Sigma), 5 µg/ml aprotinin, 5 µg/ml leupeptin, 1 µg/ml pepstatin A, and 2 mM PMSF. After detergent solubilization, binding with anti-GFP nanobody beads, and washing with Buffer W1 as described above, ATP13A2 was eluted by incubating the beads with ∼10 ug/mL 3C protease in a wash buffer (Buffer W2) containing 25 mM Tris pH 7.5, 100 mM NaCl, 1 mM EDTA, 1 mM DTT, 10 mM MgCl_2_, 0.1 mM SPM, 0.04% DDM, 0.0048% CHS, 0.0016% dipalmitoyl PA (DPPA; Echelon), and 0.001% phosphatidylinositol 3,5-bisphosphate diC16 (PI(3,5)P2-diC16; Echelon) for ∼14 h. The eluate was concentrated and injected into a Superose 6 Increase column equilibrated with a running buffer (Buffer W3) containing 25 mM Tris pH 7.5, 100 mM NaCl, 1 mM EDTA, 1 mM DTT, 5 mM MgCl_2_, 0.04% DDM, 0.0048% CHS, and 0.0016% DPPA. Purified and concentrated ATP13A2 was incubated with 1 mM AMP-PCP for 1 h at 4°C before grid preparation. We note that ATP13A2 prepared in this way was found to be in the E2-Pi state and failed to bind AMP-PCP.

For WT ATP13A2 prepared with BeF_3_ (sample for maps 3 and 4) and AlF_4_ (sample for map 5), the phosphate analogs (2 mM BeF_3_ or 2 mM AlF_4_) and 10 mM MgCl_2_ were added during detergent solubilization of the crude membrane pellet and were maintained in all subsequent buffers during the purification. For the AlF_4_ purification, inhibitor concentration was reduced to 1 mM AlF_4_ and 5 mM MgCl_2_ during size-exclusion chromatography.

For WT ATP13A2 prepared with ADP (sample for maps 6 and 7), 2 mM ADP and 5 mM MgCl_2_ was added during detergent solubilization of the crude membrane pellet and was maintained in all subsequent buffers during the purification. In addition, 2 mM AlF_4_ was added to the running buffer for size-exclusion chromatography.

For the D508N mutant bound to ATP (sample for map 8), ATP13A2 was purified according to the base procedure and incubated with 2 mM ATP and 5 mM MgCl_2_ for 1 h at 4°C before grid preparation.

The D458N/D962N mutant in its apo form (sample for maps 9 and 10) was purified according to the base procedure.

For the D458N/D962N mutant bound to AlF_4_ (sample for map 11), 2 mM AlF_4_ and 10 mM MgCl_2_ were added during detergent solubilization of the crude membrane pellet and maintained in all subsequent buffers during the purification, except during size-exclusion chromatography where the inhibitor concentration was reduced to 1 mM AlF_4_ and 5 mM MgCl_2_ in the running buffer.

### Cryo-EM grid preparation and data acquisition

To prepare cryo-EM grids, 3 μL of the ATP13A2 sample were applied to a glow-discharged (PELCO easiGlow; 0.39 mBar, 25–30 mA, 40–45 s) gold holey carbon grid (Quantifoil R 1.2/1.3, 400 mesh). The grid was blotted for 3–4 s and plunge-frozen in liquid-nitrogen-cooled liquid ethane using Vitrobot Mark IV (FEI) operated at 4° C and 100% humidity. Whatman No. 1 filter paper was used to blot the samples.

The WT-BeF_3_, WT-AlF_4_, WT-AMP-PCP, and D508N-ATP datasets were collected on a Titan Krios G2 electron microscope (FEI), operated at an acceleration voltage of 300 kV and equipped with a Gatan Quantum Image Filter (slit width of 20 eV). The WT-ADP-AlF4 and the D458N/D962N-apo datasets were collected on a Titan Krios G3i electron microscope (FEI), operated at an acceleration voltage of 300 kV and equipped with a Gatan Quantum Image Filter (slit width of 20 eV). The D458N/D962N-AlF_4_ dataset was collected on a Talos Arctica electron microscope (FEI), operated at an acceleration voltage of 200 kV. Dose-fractionated images (∼50 electrons per Å^2^ applied over 42 or 50 frames) were recorded on a K3 direct electron detector (Gatan) using the super-resolution mode. All datasets were collected using SerialEM (Mastronarde, 2005) with image-beam-shift multiple recording (typically acquiring 9 movies per stage movement with one movie per hole). Coma induced by image-beam shift was corrected by beam tilt compensation in SerialEM. The physical pixel size was 0.911 Å for the Krios G2 datasets, 1.049 Å for the Krios G3i datasets, and 1.115 Å for the Arctica dataset. Target defocus values were typically set from −0.8 to –2.0 μm.

### Cryo-EM structural determination

Movies were initially preprocessed using Warp (Tegunov and Cramer, 2019), by motion-correcting and estimating contrast transfer function (CTF) and defocus parameters with 7-by-5 tiling. Micrographs were manually inspected to remove micrographs that were not suitable for image analysis, largely those containing crystalline ice. Particles were automatically picked by Warp and were extracted with a box size of 320 pixels for the WT-apo, WT-BeF_3_, WT-AlF_4_, WT-AMP-PCP, and D508N-ATP datasets or 300 pixels for the WT-ADP-AlF_4_, D458N/D962N-apo, and D458N/D962N-AlF_4_ datasets. All subsequent image processing was performed in cryoSPARC v2 (Punjani et al., 2017), as described in detail below. Except for the WT-apo, D458N/D962N-apo, and D458N/D962N-AlF_4_ datasets where the final maps were reconstructed from particles picked in Warp, movies were reprocessed and particles were repicked in cryoSPARC. The particles selected from Warp were used to generate initial maps in cryoSPARC, which were subsequently used as a template for the second round of particle picking and heterogenous refinement in cryoSPARC.

#### (1) WT ATP13A2 prepared without any nucleotide or phosphate analog (map 1)

605,982 particles were automatically picked in Warp from 2,339 movies. Particles were imported into cryoSPARC for reference-free 2D classification, and classes without clear protein features (mostly empty micelles) were removed. 357,815 particles selected from the 2D classification were subjected to ab initio reconstruction, yielding three initial models (Classes 1 to 3). Clear features of ATP13A2 appeared in one of these classes (Class 1). The particles (357,815 particles) were classified by three rounds of heterogeneous refinement using the ab initio reconstructions. The final set of 164,606 particles was used for 3D reconstruction by non-uniform (NU) refinement and local and global CTF refinement to yield a map at 3.7-Å resolution.

#### (2) WT ATP13A2 prepared with AMP-PCP (map 2)

The initial set of 670,039 particles automatically picked in Warp from 2,374 movies was used for 2D classification. The 230,844 particles selected from the 2D classification (excluding mainly empty micelles) were subjected to ab initio reconstruction, yielding three initial models (Classes 1 to 3). Clear features of ATP13A2 appeared in one of these classes (Class 2). The particles (230,844 particles) selected from the 2D classification were classified by a round of heterogeneous refinement using the ab initio reconstructions. The resulting particles (146,852 particles) were used for 3D reconstruction by NU refinement, yielding a map at 3.7-Å resolution for the E2-Pi conformation. Raw movies (2,374 movies) were then imported into cryoSPARC for patch-based motion correction (2x pixel-binned after motion correction) and CTF estimation. 2,346 micrographs were selected, and a total of 1,001,323 particles were picked with lowpass filtered templates generated from the 3.7-Å-resolution 3D reconstruction of the E2-Pi state. Particles were extracted with a box size of 320 pixels, Fourier-cropped to 160 pixels, and subjected to a round of 2D classification. Selected particles from 2D classification (297,969 particles) were subjected to a round of heterogeneous refinement with the three initial models generated from the Warp particles. 173,886 particles were classified into Class 2 showing ATP13A3 features, and this final set of particles was subjected to NU refinement and local and global CTF refinements to yield a map at 3.2-Å resolution.

#### (3) WT ATP13A2 prepared with BeF_3_^−^ (maps 3 and 4)

A summary of the single particle-analysis is outlined in Fig. S3A. The analysis was processed similarly to WT ATP13A2 prepared with AMP-PCP. The initial set of 1,038,661 particles automatically picked in Warp from 3,431 movies was used for 2D classification. A subset (195,032 particles) of the 675,085 particles selected from the 2D classification was subjected to ab initio reconstruction, yielding three initial models (Classes 1 to 3). Clear features of ATP13A2 appeared in two structurally distinct classes (Classes 2 and 3). The selected particles (675,085 particles) were classified by a round of heterogeneous refinement using the ab initio reconstructions, where Class 1 was duplicated to aid the removal of poor-quality particles (i.e., Classes 1a, 1b, 2, and 3). Particles in Class 2 (307,364 particles) were subjected to an additional round of heterogeneous refinement. The resulting particles (281,232 particles) were used for 3D reconstruction by NU refinement and local CTF refinement, yielding a map at 3.0-Å resolution for the E2P-like conformation. Particles in Class 3 (224,085 particles) were separately subjected to an additional round of heterogeneous refinement. The resulting particles (205,001 particles) were used for 3D reconstruction by NU refinement, yielding a map at 3.6-Å resolution for the E2-Pi conformation. Raw movies (3,431 movies) were then imported into cryoSPARC for patch-based motion correction and CTF estimation. 3,395 micrographs were selected, and a total of 1,437,359 particles were picked with 2D templates generated from the 3.0-Å-resolution 3D reconstruction of the E2P-like state. Particles were extracted with a box size of 320 pixels, Fourier-cropped to 160 pixels, and subjected to2D classification. Selected particles from 2D classification (873,004 particles) were then subjected to heterogeneous refinement with the three initial models generated from the Warp particles.

Among 427,905 particles that were classified into Class 2 (E2P-like structure), a subset (200,000 particles) was subjected to a round of ab initio reconstruction to yield three maps. Two additional rounds of heterogeneous refinement (starting from all 427,905 particles) were performed to remove poor-quality particles. The final set of 386,118 particles was subjected to NU refinement and local and global CTF refinements to yield a map at 2.8-Å resolution (map 3).

311,737 particles were classified into Class 3 (E2-Pi structure) and subjected to the essentially same 3D classification procedure as described for map 3. This generated a set of 266,167 particles, which was further narrowed to 253,411 particles using class probability filtering. This final set of 253,411 particles was subjected to NU refinement and local and global CTF refinements to yield a map at 3.1-Å resolution (map 4).

#### (4) WT ATP13A2 prepared with AlF_4_ (map 5)

The analysis was processed similarly to WT ATP13A2 prepared with AMP-PCP. The initial set of 1,152,292 particles automatically picked in Warp from 3,519 movies was used for 2D classification, and 695,816 particles selected from the 2D classification were subjected to ab initio reconstruction, yielding three initial models (Classes 1 to 3). Clear features of ATP13A2 appeared in two of these classes (Classes 2 and 3), which were structurally similar. The particles (695,816 particles) were classified by two rounds of heterogeneous refinement using the ab initio reconstructions. The resulting particles (512,952 particles) were used for 3D reconstruction by NU refinement, yielding a map at 3.0-Å resolution. Raw movies (3,519 movies) were then imported into cryoSPARC for patch-based motion correction and CTF estimation. 3,420 movies were selected, and a total of 1,716,479 particles were picked with templates generated from the 3.1-Å-resolution 3D reconstruction. Particles were extracted with a box size of 320 pixels, Fourier-cropped to 160 pixels, and subjected to 2D classification. Selected particles from 2D classification (946,867 particles) were then subjected to heterogeneous refinement with the initial models generated from the Warp particles. 679,631 particles were classified into a class showing ATP13A3 features, and another round of ab initio reconstruction, heterogeneous refinement, and class probability filtering was used to further remove poor-quality particles. The final set of 596,369 particles was subjected to NU refinement and local and global CTF refinements to yield a map at 2.8-Å resolution.

#### (5) WT ATP13A2 prepared with ADP (maps 6 and 7)

A summary of the single particle-analysis is outlined in Fig. S3B. The analysis was processed similarly to WT ATP13A2 prepared with AMP-PCP. The initial set of 1,159,048 particles automatically picked in Warp from 3,361 movies was used for 2D classification. A subset (257,440 particles) of the 746,647 particles selected from the 2D classification were subjected to ab initio reconstruction, yielding three initial models. Clear features of ATP13A2 appeared in two of these classes (Classes 1 and 2), which were structurally distinct. These particles (746,647 particles) were classified by a round of heterogeneous refinement using the ab initio reconstructions. Particles in Class 1 (490,426 particles) were used for 3D reconstruction by NU refinement, yielding a map at 2.5-Å resolution for the E2-Pi-like conformation. Particles in Class 2 (150,444 particles) were separately used for 3D reconstruction by NU refinement, yielding a map at 3.0-Å resolution for the E1P-ADP-like conformation. Raw movies (3,361 movies) were then imported into cryoSPARC for patch-based motion correction and CTF estimation. 3,347 micrographs were selected, and a total of 1,311,023 particles were picked with templates generated from the 2.5-Å-resolution 3D reconstruction of the E2-Pi-like state. Particles were extracted with a box size of 300 pixels, Fourier-cropped to 150 pixels, and subjected to 2D classification. Selected particles from 2D classification (892,802 particles) were then subjected to heterogeneous refinement with the four initial models [Class 1 (E2-Pi-like), Class 2 (E1P-ADP-like), Class 3a (poor-quality class), Class 3b (poor-quality class)] generated from the Warp particles.

570,133 particles that were classified into Class 1 were subjected to another round of heterogeneous refinement to remove junk particles. The final set of 462,490 particles was subjected to NU refinement and local and global CTF refinements to yield a map at 2.5-Å resolution (map 6). 184,257 particles that were classified into Class 2 were subjected to another round of ab initio reconstruction and heterogeneous refinement to remove junk particles. The final set of 163,433 particles was subjected to NU refinement and local and global CTF refinements to yield a map at 2.9-Å resolution (map 7).

#### (6) D508N mutant prepared with ATP (map 8)

The analysis was processed similarly to WT ATP13A2 prepared with AMP-PCP. An initial set of 953,226 particles was automatically picked in Warp from 2,833 movies, and a subset of these particles (399,085 particles) were used for 2D classification 247,353 particles were selected from the 2D classification and subjected to ab initio reconstruction, yielding three initial models (Classes 1 to 3). Clear features of ATP13A2 appeared in one of these classes (Class 1). The particles (247,353 particles) were classified by a round of heterogeneous refinement using the ab initio reconstructions. The resulting particles (139,467 particles) were used for 3D reconstruction by NU refinement, yielding a map at 3.5-Å resolution for the E1-ATP-like conformation. Raw movies (2,833 movies) were then imported into cryoSPARC for patch-based motion correction and CTF estimation. 2,755 micrographs were selected, and a total of 1,431,882 particles were picked with templates generated from the 3.5-Å-resolution 3D reconstruction of the E1-ATP-like state. Particles were extracted with a box size of 320 pixels, Fourier-cropped to 160 pixels, and subjected to two rounds of 2D classification. Selected particles from 2D classification (756,595 particles) were subjected to a round of heterogeneous refinement with the three initial models [Class 1 (E1-ATP-like), Class 2 (poor-quality class), and Class 3 (poor-quality class)] generated from the Warp particles. 459,541 particles were classified into Class 1, and this final set of particles was subjected to NU refinement and local and global CTF refinements to yield a map at 2.8-Å resolution.

#### (7) D458N/D962N mutant in its apo form (maps 9 and 10)

The initial set of 1,181,866 particles automatically picked in Warp was used for 2D classification. 589,548 particles selected from the 2D classification were subjected to ab initio reconstruction, yielding three initial models (Classes 1 to 3). Clear features of ATP13A2 appeared in one of these classes. The particles (589,548 particles) were classified by two rounds of heterogeneous refinement using the ab initio reconstructions. The resulting particles (375,210 particles) were subjected to a final round of heterogeneous refinement to separate two different conformations of the N domain (Classes 1a and 1b). Particles in Class 1b (186,331 particles) were used for 3D reconstruction by NU refinement and local CTF refinement, yielding a map at 2.9-Å resolution for the first E1 (apo) conformation (map 9). Particles in Class 1b (188,879 particles) were separately used for three-dimensional (3D) reconstruction by NU refinement and local CTF refinement, yielding a map at 2.9-Å resolution for the second E1 (apo) conformation (map 10).

#### (8) D458N/D962N mutant prepared with AlF_4_ (map 11)

The initial set of 1,406,963 particles automatically picked in Warp from 3,214 movies was used for 2D classification. 746,533 particles selected from the 2D classification were subjected to ab initio reconstruction, yielding four initial models (Classes 1 to 4). Clear features of ATP13A2 appeared in two of these classes (Classes 1 and 2), which were structurally similar. The particles (746,533 particles) were classified by two rounds of heterogeneous refinement and an additional round of ab initio reconstruction. The final set of 377,365 particles (from Class “1+2”) was subjected to NU refinement and local and global CTF refinements to yield a map at 3.2-Å resolution (map 11).

#### (9) Combined post-hydrolysis E2-Pi structure of WT ATP13A2 (map 12)

To improve the quality of the post-hydrolysis E2-Pi state map, particles used to generate three maps (164,606 particles from map 1, 173,886 particles from map 2, and 253,411 particles from map 4) were combined (a total of 591,903 particles) and subjected to NU refinement to yield a map at 3.0-Å resolution.

### Atomic Model Building

The initial atomic model was built *de novo* into the sharpened map of WT-BeF_3_ structure using Coot (Emsley et al., 2010). Models for the other structures were built after rigid-body fitting of individual domains into the corresponding maps using the model of the WT-BeF_3_ structure and rounds of local refinement in Coot. The following sequences were not modeled because they were poorly resolved in the density maps: N to 33, 108 to 156 (a part of the NTD; residues 108 to 110 are resolved in E1P model), 409 to 415 (connection between the A-domain and TM3), 582 to 586 (a loop in the N-domain), 595 to 613 (a part of the N-domain), and 1169 to 1175 (C-terminus). Residue side chains with poor density were truncated at the β-carbon. Model refinement was performed using Phenix (*phenix*.*real_space_refine*) (Afonine et al., 2018) with secondary structure restraints and with the refinement resolution limit set to the overall resolution of the map. For E1 (apo), E1P, and E2-Pi structures, the N domain was treated as a rigid body by applying a strong reference model restraint generated from the N-domain of the refined WT-BeF_3_^−^ structure, because cryo-EM density features were relatively poor in this region. Structural validation was performed using MolProbity (Chen et al., 2010) in the Phenix package. Protein electrostatics were calculated using the Adaptive Poisson-Boltzmann Solver (Baker et al., 2001; Dolinsky et al., 2004) with default parameters built in PyMOL (with monovalent ion concentrations of 0.15 M each). Substrate-transport pathways were detected by Caver 3.0 (Chovancova et al., 2012). UCSF Chimera (Pettersen et al., 2004), Chimera X (Goddard et al., 2018), and PyMOL (Schrödinger) were used to prepare structural figures in the paper.

### Coarse-grained simulations of ATP13A2 in a lysosomal membrane

Coarse-grained molecular dynamics simulations of ATP13A2 were performed in a model lysosomal membrane using the Martini 2.2 force field (de Jong et al., 2013). Coarse-grained Martini structures of ATP13A2 E2-P and E2-Pi were prepared using the software *martinize*.*py* (de Jong et al., 2013). The protein internal structure was constrained with an elastic network with a bond force constant of 200 kJ/mol/nm^2^, and lower and upper bond cut-offs of 0.5 nm and 0.9 nm, respectively.

The ATP13A2 model (E2P or E2-Pi) was positioned in an asymmetric lipid bilayer with one of two lipid compositions, *u* (unphosphorylated) and *p* (phosphorylated), using the program *insane*.*py* (Wassenaar et al., 2015). The bilayer compositions mimic a lysosomal membrane with different cytosolic (upper) and luminal (lower) leaflets as described for the plasma membrane (Lorent et al., 2020; van Meer et al., 2008). The *u* composition consists of 15.2% cholesterol, 19.8% phosphatidylcholine (13.2% PIPC and 6.6% POPC), 32.0% phosphatidylethanolamine (DOPE), 23.0% phosphatidic acid (DOPA), 3.0% phosphatidylserine (PAPS), and 7.0% phosphatidylinositol (PAPI) in the cytosolic leaflet, and 38.8% cholesterol, 30.8% PIPC, 15.4% POPC, and 15.0% sphingomyelin (DPSM) in the lumenal leaflet. DOPA was used as substituent for the abundant lysosomal lipid bis(monoacylglyceryl)phosphate, having the same charge and identical acyl chain length and saturation. The *p* composition is the same as the *u* composition except that a half of PAPI lipids (i.e., 3.5%) were replaced by PI(3,5)P_2_. We defined parameters for the non-standard lipid POPI(3,5)P_2_ following the procedure described previously (Lopez et al., 2013). The system was solvated with water beads using *insane*.*py*.

In the four simulation systems E2P/*u*, E2P/*p*, E2-Pi/*u*, and E2-Pi/*p*, we combined the indicated protein conformations and lipid bilayer compositions. All systems had initial box edge lengths of 25 nm in *x/y*-direction and 16 nm in z-direction. Gromacs 2020/3 (Abraham et al., 2015) was used to preprocess and simulate the four systems. 10% of the water beads were replaced by ‘‘anti-freeze’’ (WF) particles. The systems were neutralized with sodium ion beads. The energy was minimized by steepest descent up to convergence (< 1000.0 kJ/mol/nm between adjacent steps). We maintained a temperature of 300 K with the v-rescale thermostat (Bussi et al., 2007). We equilibrated the systems first in the *NVT* ensemble for 100 ns, then in the *NPT* ensemble for 100 ns using the Berendsen barostat (Berendsen et al., 1984) at 1 bar with semiisotropic pressure coupling and τ_c_=5 ps. Production runs were carried out in the *NPT* ensemble for 10 µs using the Parrinello-Rahman barostat (Parrinello and Rahman, 1981) at 1 bar with τ_c_ of 12 ps. Simulation structures were recorded every 1.5 ns. The lipid structure around the proteins was analyzed with custom *python* scripts using the packages *numpy* (www.numpy.org) and *MDAnalysis* (Gowers et al., 2016). A minimum bead distance of <0.6 nm was used to define lipid-protein contacts. Plots were created with the *python* package *matplotlib* (www.matplotlib.org).

## Data availability

Cryo-EM density maps and atomic models are available under accession codes EMDB XXXXX-YYYYYY and PDB XXXX-YYYY.

## Notes

### Competing Interest Statement

The authors have declared no competing interest.

## References

Abbas, M.M., Govindappa, S.T., Sheerin, U.M., Bhatia, K.P., and Muthane, U.B. (2017). Exome Sequencing Identifies a Novel Homozygous Missense ATP13A2 Mutation. Mov Disord Clin Pract 4, 132– 135.

Abraham, M.J., Murtola, T., Schulz, R., Páll, S., Smith, J.C., Hess, B., and Lindahl, E. (2015). GROMACS: High performance molecular simulations through multi-level parallelism from laptops to supercomputers. SoftwareX 1-2, 19–25.

Afonine, P.V., Poon, B.K., Read, R.J., Sobolev, O.V., Terwilliger, T.C., Urzhumtsev, A., and Adams, P.D. (2018). Real-space refinement in PHENIX for cryo-EM and crystallography. Acta Crystallogr D Struct Biol 74, 531–544.

Aouida, M., Leduc, A., Poulin, R., and Ramotar, D. (2005). AGP2 encodes the major permease for high affinity polyamine import in Saccharomyces cerevisiae. The Journal of biological chemistry 280, 24267– 24276.

Bai, L., Kovach, A., You, Q., Hsu, H.-C., Zhao, G., and Li, H. (2019). Autoinhibition and activation mechanisms of the eukaryotic lipid flippase Drs2p-Cdc50p. Nature Communications 10, 4142.

Bai, L., You, Q., Jain, B.K., Duan, H.D., Kovach, A., Graham, T.R., and Li, H. (2020). Transport mechanism of P4 ATPase phosphatidylcholine flippases. eLife 9, e62163.

Baker, N.A., Sept, D., Joseph, S., Holst, M.J., and McCammon, J.A. (2001). Electrostatics of nanosystems: application to microtubules and the ribosome. Proc Natl Acad Sci U S A 98, 10037–10041.

Berendsen, H.J.C., Postma, J.P.M., van Gunsteren, W.F., DiNola, A., and Haak, J.R. (1984). Molecular dynamics with coupling to an external bath. The Journal of Chemical Physics 81, 3684–3690.

Bras, J., Verloes, A., Schneider, S.A., Mole, S.E., and Guerreiro, R.J. (2012). Mutation of the parkinsonism gene ATP13A2 causes neuronal ceroid-lipofuscinosis. Hum Mol Genet 21, 2646–2650.

Bublitz, M., Poulsen, H., Morth, J.P., and Nissen, P. (2010). In and out of the cation pumps: P-Type ATPase structure revisited. Current Opinion in Structural Biology 20, 431–439.

Bussi, G., Donadio, D., and Parrinello, M. (2007). Canonical sampling through velocity rescaling. J Chem Phys 126, 014101.

Chen, C.M., Lin, C.H., Juan, H.F., Hu, F.J., Hsiao, Y.C., Chang, H.Y., Chao, C.Y., Chen, I.C., Lee, L.C., Wang, T.W., et al. (2011). ATP13A2 variability in Taiwanese Parkinson’s disease. Am J Med Genet B Neuropsychiatr Genet 156B, 720–729.

Chen, V.B., Arendall, W.B., 3rd, Headd, J.J., Keedy, D.A., Immormino, R.M., Kapral, G.J., Murray, L.W., Richardson, J.S., and Richardson, D.C. (2010). MolProbity: all-atom structure validation for macromolecular crystallography. Acta Crystallogr D Biol Crystallogr 66, 12–21.

Chovancova, E., Pavelka, A., Benes, P., Strnad, O., Brezovsky, J., Kozlikova, B., Gora, A., Sustr, V., Klvana, M., Medek, P., et al. (2012). CAVER 3.0: a tool for the analysis of transport pathways in dynamic protein structures. PLoS Comput Biol 8, e1002708.

Clausen, M.V., Hilbers, F., and Poulsen, H. (2017). The Structure and Function of the Na,K-ATPase Isoforms in Health and Disease. Front Physiol 8, 371.

de Jong, D.H., Singh, G., Bennett, W.F., Arnarez, C., Wassenaar, T.A., Schafer, L.V., Periole, X., Tieleman, D.P., and Marrink, S.J. (2013). Improved Parameters for the Martini Coarse-Grained Protein Force Field. J Chem Theory Comput 9, 687–697.

De La Hera, D.P., Corradi, G.R., Adamo, H.P., and De Tezanos Pinto, F. (2013). Parkinson’s disease-associated human P5B-ATPase ATP13A2 increases spermidine uptake. Biochem J 450, 47–53.

Dehay, B., Ramirez, A., Martinez-Vicente, M., Perier, C., Canron, M.-H., Doudnikoff, E., Vital, A., Vila, M., Klein, C., and Bezard, E. (2012). Loss of P-type ATPase ATP13A2/PARK9 function induces general lysosomal deficiency and leads to Parkinson disease neurodegeneration. Proceedings of the National Academy of Sciences 109, 9611–9616.

Di Fonzo, A., Chien, H.F., Socal, M., Giraudo, S., Tassorelli, C., Iliceto, G., Fabbrini, G., Marconi, R., Fincati, E., Abbruzzese, G., et al. (2007). ATP13A2 missense mutations in juvenile parkinsonism and young onset Parkinson disease. Neurology 68, 1557–1562.

Djarmati, A., Hagenah, J., Reetz, K., Winkler, S., Behrens, M.I., Pawlack, H., Lohmann, K., Ramirez, A., Tadic, V., Bruggemann, N., et al. (2009). ATP13A2 variants in early-onset Parkinson’s disease patients and controls. Mov Disord 24, 2104–2111.

Dolinsky, T.J., Nielsen, J.E., McCammon, J.A., and Baker, N.A. (2004). PDB2PQR: an automated pipeline for the setup of Poisson-Boltzmann electrostatics calculations. Nucleic Acids Res 32, W665–667.

Dyla, M., Basse Hansen, S., Nissen, P., and Kjaergaard, M. (2019). Structural dynamics of P-type ATPase ion pumps. Biochemical Society Transactions 47, 1247–1257.

Emsley, P., Lohkamp, B., Scott, W.G., and Cowtan, K. (2010). Features and development of Coot. Acta Crystallogr D Biol Crystallogr 66, 486–501.

Estiar, M.A., Leveille, E., Spiegelman, D., Dupre, N., Trempe, J.F., Rouleau, G.A., and Gan-Or, Z. (2020). Clinical and genetic analysis of ATP13A2 in hereditary spastic paraplegia expands the phenotype. Mol Genet Genomic Med 8, e1052.

Estrada-Cuzcano, A., Martin, S., Chamova, T., Synofzik, M., Timmann, D., Holemans, T., Andreeva, A., Reichbauer, J., De Rycke, R., Chang, D.-I., et al. (2017). Loss-of-function mutations in the ATP13A2/ PARK9 gene cause complicated hereditary spastic paraplegia (SPG78). Brain 140, 287–305.

Exner, N., Lutz, A.K., Haass, C., and Winklhofer, K.F. (2012). Mitochondrial dysfunction in Parkinson’s disease: molecular mechanisms and pathophysiological consequences. EMBO J 31, 3038–3062.

Feng, Z., Zhao, Y., Li, T., Nie, W., Yang, X., Wang, X., Wu, J., Liao, J., and Zou, Y. (2020). CATP-8/P5A ATPase Regulates ER Processing of the DMA-1 Receptor for Dendritic Branching. Cell Reports 32, 108101.

Fujita, M., Fujita, Y., Iuchi, S., Yamada, K., Kobayashi, Y., Urano, K., Kobayashi, M., Yamaguchi-Shinozaki, K., and Shinozaki, K. (2012). Natural variation in a polyamine transporter determines paraquat tolerance in Arabidopsis. Proceedings of the National Academy of Sciences of the United States of America 109, 6343–6347.

Gitler, A.D., Chesi, A., Geddie, M.L., Strathearn, K.E., Hamamichi, S., Hill, K.J., Caldwell, K.A., Caldwell, G.A., Cooper, A.A., Rochet, J.-C., et al. (2009). α-Synuclein is part of a diverse and highly conserved interaction network that includes PARK9 and manganese toxicity. Nature Genetics 41, 308–315.

Goddard, T.D., Huang, C.C., Meng, E.C., Pettersen, E.F., Couch, G.S., Morris, J.H., and Ferrin, T.E. (2018). UCSF ChimeraX: Meeting modern challenges in visualization and analysis. Protein Sci 27, 14–25.

Gowers, R.J., Linke, J., Barnoud, J., Reddy, T.J.E., Melo, M.N., Seyler, S.L., Dotson, D.L., Domanski, J., Buchoux, S., Kenney, I.M., et al. (2016). MDAnalysis: A Python package for the rapid analysis of molecular dynamics simulations. Proceedings of the 15th Python in Science Conference, SciPy, 98–105.

Grunewald, A., Arns, B., Seibler, P., Rakovic, A., Munchau, A., Ramirez, A., Sue, C.M., and Klein, C. (2012). ATP13A2 mutations impair mitochondrial function in fibroblasts from patients with Kufor-Rakeb syndrome. Neurobiol Aging 33, 1843 e1841–1847.

Hamouda, N.N., Van den Haute, C., Vanhoutte, R., Sannerud, R., Azfar, M., Mayer, R., Cortes Calabuig, A., Swinnen, J.V., Agostinis, P., Baekelandt, V., et al. (2020). ATP13A3 is a major component of the enigmatic mammalian polyamine transport system. J Biol Chem.

Handa, A.K., Fatima, T., and Mattoo, A.K. (2018). Polyamines: Bio-Molecules with Diverse Functions in Plant and Human Health and Disease. Front Chem 6, 10.

Heinick, A., Urban, K., Roth, S., Spies, D., Nunes, F., Phanstiel, O.t., Liebau, E., and Luersen, K. (2010). Caenorhabditis elegans P5B-type ATPase CATP-5 operates in polyamine transport and is crucial for norspermidine-mediated suppression of RNA interference. FASEB J 24, 206–217.

Hiasa, M., Miyaji, T., Haruna, Y., Takeuchi, T., Harada, Y., Moriyama, S., Yamamoto, A., Omote, H., and Moriyama, Y. (2014). Identification of a mammalian vesicular polyamine transporter. In Sci Rep, pp. 6836.

Hiraizumi, M., Yamashita, K., Nishizawa, T., and Nureki, O. (2019). Cryo-EM structures capture the transport cycle of the P4-ATPase flippase. Science 365, 1149–1155.

Holemans, T., Sørensen, D.M., van Veen, S., Martin, S., Hermans, D., Kemmer, G.C., Van den Haute, C., Baekelandt, V., Günther Pomorski, T., Agostinis, P., et al. (2015). A lipid switch unlocks Parkinson’s disease-associated ATP13A2. Proceedings of the National Academy of Sciences 112, 9040–9045.

Igarashi, K., and Kashiwagi, K. (1999). Polyamine transport in bacteria and yeast. The Biochemical journal 344 Pt 3, 633–642.

Igarashi, K., and Kashiwagi, K. (2010). Modulation of cellular function by polyamines. The International Journal of Biochemistry & Cell Biology 42, 39–51.

Kashiwagi, K., Pistocchi, R., Shibuya, S., Sugiyama, S., Morikawa, K., and Igarashi, K. (1996). Spermidine-preferential uptake system in Escherichia coli. Identification of amino acids involved in polyamine binding in PotD protein. J Biol Chem 271, 12205–12208.

Kuhlbrandt, W. (2004). Biology, structure and mechanism of P-type ATPases. Nat Rev Mol Cell Biol 5, 282–295.

López-Marqués, R.L., Gourdon, P., Günther Pomorski, T., and Palmgren, M. (2020). The transport mechanism of P4 ATPase lipid flippases. Biochemical Journal 477, 3769–3790.

Lopez, C.A., Sovova, Z., van Eerden, F.J., de Vries, A.H., and Marrink, S.J. (2013). Martini Force Field Parameters for Glycolipids. J Chem Theory Comput 9, 1694–1708.

Lorent, J.H., Levental, K.R., Ganesan, L., Rivera-Longsworth, G., Sezgin, E., Doktorova, M., Lyman, E., and Levental, I. (2020). Plasma membranes are asymmetric in lipid unsaturation, packing and protein shape. Nat Chem Biol 16, 644–652.

Mastronarde, D.N. (2005). Automated electron microscope tomography using robust prediction of specimen movements. J Struct Biol 152, 36–51.

McKenna, M.J., Sim, S.I., Ordureau, A., Wei, L., Harper, J.W., Shao, S., and Park, E. (2020). The endoplasmic reticulum P5A-ATPase is a transmembrane helix dislocase. Science 369, eabc5809.

McNeil-Gauthier, A.L., Brais, B., Rouleau, G., Anoja, N., and Ducharme, S. (2019). Successful treatment of psychosis in a patient with Kufor-Rakeb syndrome with low dose aripiprazole: a case report. Neurocase 25, 133–137.

Møller, A.B., Asp, T., Holm, P.B., and Palmgren, M.G. (2008). Phylogenetic analysis of P5 P-type ATPases, a eukaryotic lineage of secretory pathway pumps. Molecular Phylogenetics and Evolution 46, 619–634.

Morth, J.P., Pedersen, B.P., Toustrup-Jensen, M.S., Sørensen, T.L.-M., Petersen, J., Andersen, J.P., Vilsen, B., and Nissen, P. (2007). Crystal structure of the sodium–potassium pump. Nature 450, 1043–1049.

Nakanishi, H., Nishizawa, T., Segawa, K., Nureki, O., Fujiyoshi, Y., Nagata, S., and Abe, K. (2020). Transport Cycle of Plasma Membrane Flippase ATP11C by Cryo-EM. Cell Reports 32, 108208.

Ning, Y.P., Kanai, K., Tomiyama, H., Li, Y., Funayama, M., Yoshino, H., Sato, S., Asahina, M., Kuwabara, S., Takeda, A., et al. (2008). PARK9-linked parkinsonism in eastern Asia: mutation detection in ATP13A2 and clinical phenotype. Neurology 70, 1491–1493.

Olesen, C., Picard, M., Winther, A.M., Gyrup, C., Morth, J.P., Oxvig, C., Moller, J.V., and Nissen, P. (2007). The structural basis of calcium transport by the calcium pump. Nature 450, 1036–1042.

Palmgren, M.G., and Nissen, P. (2011). P-Type ATPases. Annual Review of Biophysics 40, 243–266.

Park, J.S., Mehta, P., Cooper, A.A., Veivers, D., Heimbach, A., Stiller, B., Kubisch, C., Fung, V.S., Krainc, D., Mackay-Sim, A., et al. (2011). Pathogenic effects of novel mutations in the P-type ATPase ATP13A2 (PARK9) causing Kufor-Rakeb syndrome, a form of early-onset parkinsonism. Hum Mutat 32, 956–964.

Parrinello, M., and Rahman, A. (1981). Polymorphic transitions in single crystals: A new molecular dynamics method. Journal of Applied Physics 52, 7182–7190.

Pegg, A.E. (2009). Mammalian polyamine metabolism and function. IUBMB Life 61, 880–894.

Pettersen, E.F., Goddard, T.D., Huang, C.C., Couch, G.S., Greenblatt, D.M., Meng, E.C., and Ferrin, T.E. (2004). UCSF Chimera--a visualization system for exploratory research and analysis. J Comput Chem 25, 1605–1612.

Podhajska, A., Musso, A., Trancikova, A., Stafa, K., Moser, R., Sonnay, S., Glauser, L., and Moore, D.J. (2012). Common pathogenic effects of missense mutations in the P-type ATPase ATP13A2 (PARK9) associated with early-onset parkinsonism. PLoS One 7, e39942.

Punjani, A., Rubinstein, J.L., Fleet, D.J., and Brubaker, M.A. (2017). cryoSPARC: algorithms for rapid unsupervised cryo-EM structure determination. Nat Methods 14, 290–296.

Qin, Q., Zhao, T., Zou, W., Shen, K., and Wang, X. (2020). An Endoplasmic Reticulum ATPase Safeguards Endoplasmic Reticulum Identity by Removing Ectopically Localized Mitochondrial Proteins. Cell Reports 33, 108363.

Ramirez, A., Heimbach, A., Gründemann, J., Stiller, B., Hampshire, D., Cid, L.P., Goebel, I., Mubaidin, A.F., Wriekat, A.-L., Roeper, J., et al. (2006). Hereditary parkinsonism with dementia is caused by mutations in ATP13A2, encoding a lysosomal type 5 P–type ATPase. Nature Genetics 38, 1184–1191.

Ritz, C., Baty, F., Streibig, J.C., and Gerhard, D. (2015). Dose-Response Analysis Using R. PLoS One 10, e0146021.

Santoro, L., Breedveld, G.J., Manganelli, F., Iodice, R., Pisciotta, C., Nolano, M., Punzo, F., Quarantelli, M., Pappata, S., Di Fonzo, A., et al. (2011). Novel ATP13A2 (PARK9) homozygous mutation in a family with marked phenotype variability. Neurogenetics 12, 33–39.

Scholz, J., and Suppmann, S. (2017). A new single-step protocol for rapid baculovirus-driven protein production in insect cells. BMC Biotechnol 17, 83.

Shinoda, T., Ogawa, H., Cornelius, F., and Toyoshima, C. (2009). Crystal structure of the sodium-potassium pump at 2.4 A resolution. Nature 459, 446–450.

Sørensen, D.M., Buch-Pedersen, M.J., and Palmgren, M.G. (2010). Structural divergence between the two subgroups of P5 ATPases. Biochimica et Biophysica Acta (BBA) - Bioenergetics 1797, 846–855.

Sorensen, D.M., Holemans, T., van Veen, S., Martin, S., Arslan, T., Haagendahl, I.W., Holen, H.W., Hamouda, N.N., Eggermont, J., Palmgren, M., et al. (2018). Parkinson disease related ATP13A2 evolved early in animal evolution. PLoS One 13, e0193228.

Sorensen, T.L., Moller, J.V., and Nissen, P. (2004). Phosphoryl transfer and calcium ion occlusion in the calcium pump. Science 304, 1672–1675.

Spataro, R., Kousi, M., Farhan, S.M.K., Willer, J.R., Ross, J.P., Dion, P.A., Rouleau, G.A., Daly, M.J., Neale, B.M., La Bella, V., et al. (2019). Mutations in ATP13A2 (PARK9) are associated with an amyotrophic lateral sclerosis-like phenotype, implicating this locus in further phenotypic expansion. Hum Genomics 13, 19.

Sugiyama, S., Matsuo, Y., Vassylyev, D.G., Matsushima, M., Morikawa, K., Maenaka, K., Kashiwagi, K., and Igarashi, K. (1996). The 1.8-Å X-ray structure of the Escherichia coli PotD protein complexed with spermidine and the mechanism of polyamine binding. Protein Science 5, 1984–1990.

Suleiman, J., Hamwi, N., and El-Hattab, A.W. (2018). ATP13A2 novel mutations causing a rare form of juvenile-onset Parkinson disease. Brain Dev 40, 824–826.

Tegunov, D., and Cramer, P. (2019). Real-time cryo-electron microscopy data preprocessing with Warp. Nat Methods 16, 1146–1152.

Timcenko, M., Lyons, J.A., Januliene, D., Ulstrup, J.J., Dieudonné, T., Montigny, C., Ash, M.-R., Karlsen, J.L., Boesen, T., Kühlbrandt, W., et al. (2019). Structure and autoregulation of a P4-ATPase lipid flippase. Nature 571, 366–370.

Toyoshima, C., Nakasako, M., Nomura, H., and Ogawa, H. (2000). Crystal structure of the calcium pump of sarcoplasmic reticulum at 2.6 A resolution. Nature 405, 647–655.

Toyoshima, C., Nomura, H., and Tsuda, T. (2004). Lumenal gating mechanism revealed in calcium pump crystal structures with phosphate analogues. Nature 432, 361–368.

Twomey, E.C., Yelshanskaya, M.V., Vassilevski, A.A., and Sobolevsky, A.I. (2018). Mechanisms of Channel Block in Calcium-Permeable AMPA Receptors. Neuron 99, 956-968.e954.

Usenovic, M., Tresse, E., Mazzulli, J.R., Taylor, J.P., and Krainc, D. (2012). Deficiency of ATP13A2 leads to lysosomal dysfunction, α-synuclein accumulation, and neurotoxicity. J Neurosci 32.

van de Warrenburg, B.P., Schouten, M.I., de Bot, S.T., Vermeer, S., Meijer, R., Pennings, M., Gilissen, C., Willemsen, M.A., Scheffer, H., and Kamsteeg, E.J. (2016). Clinical exome sequencing for cerebellar ataxia and spastic paraplegia uncovers novel gene-disease associations and unanticipated rare disorders. Eur J Hum Genet 24, 1460–1466.

van Meer, G., Voelker, D.R., and Feigenson, G.W. (2008). Membrane lipids: where they are and how they behave. Nat Rev Mol Cell Biol 9, 112–124.

van Veen, S., Martin, S., Van den Haute, C., Benoy, V., Lyons, J., Vanhoutte, R., Kahler, J.P., Decuypere, J.- P., Gelders, G., Lambie, E., et al. (2020). ATP13A2 deficiency disrupts lysosomal polyamine export. Nature 578, 419–424.

Vrijsen, S., Besora-Casals, L., van Veen, S., Zielich, J., Van den Haute, C., Hamouda, N.N., Fischer, C., Ghesquière, B., Tournev, I., Agostinis, P., et al. (2020). ATP13A2-mediated endo-lysosomal polyamine export counters mitochondrial oxidative stress. Proceedings of the National Academy of Sciences, 201922342.

Wallings, R.L., Humble, S.W., Ward, M.E., and Wade-Martins, R. (2019). Lysosomal Dysfunction at the Centre of Parkinson’s Disease and Frontotemporal Dementia/Amyotrophic Lateral Sclerosis. Trends Neurosci 42, 899–912.

Wang, R., Tan, J., Chen, T., Han, H., Tian, R., Tan, Y., Wu, Y., Cui, J., Chen, F., Li, J., et al. (2019). ATP13A2 facilitates HDAC6 recruitment to lysosome to promote autophagosome-lysosome fusion. J Cell Biol 218, 267–284.

Wassenaar, T.A., Ingolfsson, H.I., Bockmann, R.A., Tieleman, D.P., and Marrink, S.J. (2015). Computational Lipidomics with insane: A Versatile Tool for Generating Custom Membranes for Molecular Simulations. J Chem Theory Comput 11, 2144–2155.

Williams, K. (1997). Interactions of polyamines with ion channels. Biochemical Journal 325, 289–297.

Winther, A.M., Bublitz, M., Karlsen, J.L., Moller, J.V., Hansen, J.B., Nissen, P., and Buch-Pedersen, M.J. (2013). The sarcolipin-bound calcium pump stabilizes calcium sites exposed to the cytoplasm. Nature 495, 265–269.

Yang, X., and Xu, Y. (2014). Mutations in the ATP13A2 gene and Parkinsonism: a preliminary review. Biomed Res Int 2014, 371256.

Yung, M.W., and Green, C. (1986). The binding of polyamines to phospholipid bilayers. Biochem Pharmacol 35, 4037–4041.

