## Supplemental Figures S1-7 and Tables S1-2 for "Structural basis of polyamine transport by human ATP13A2 (PARK9)"

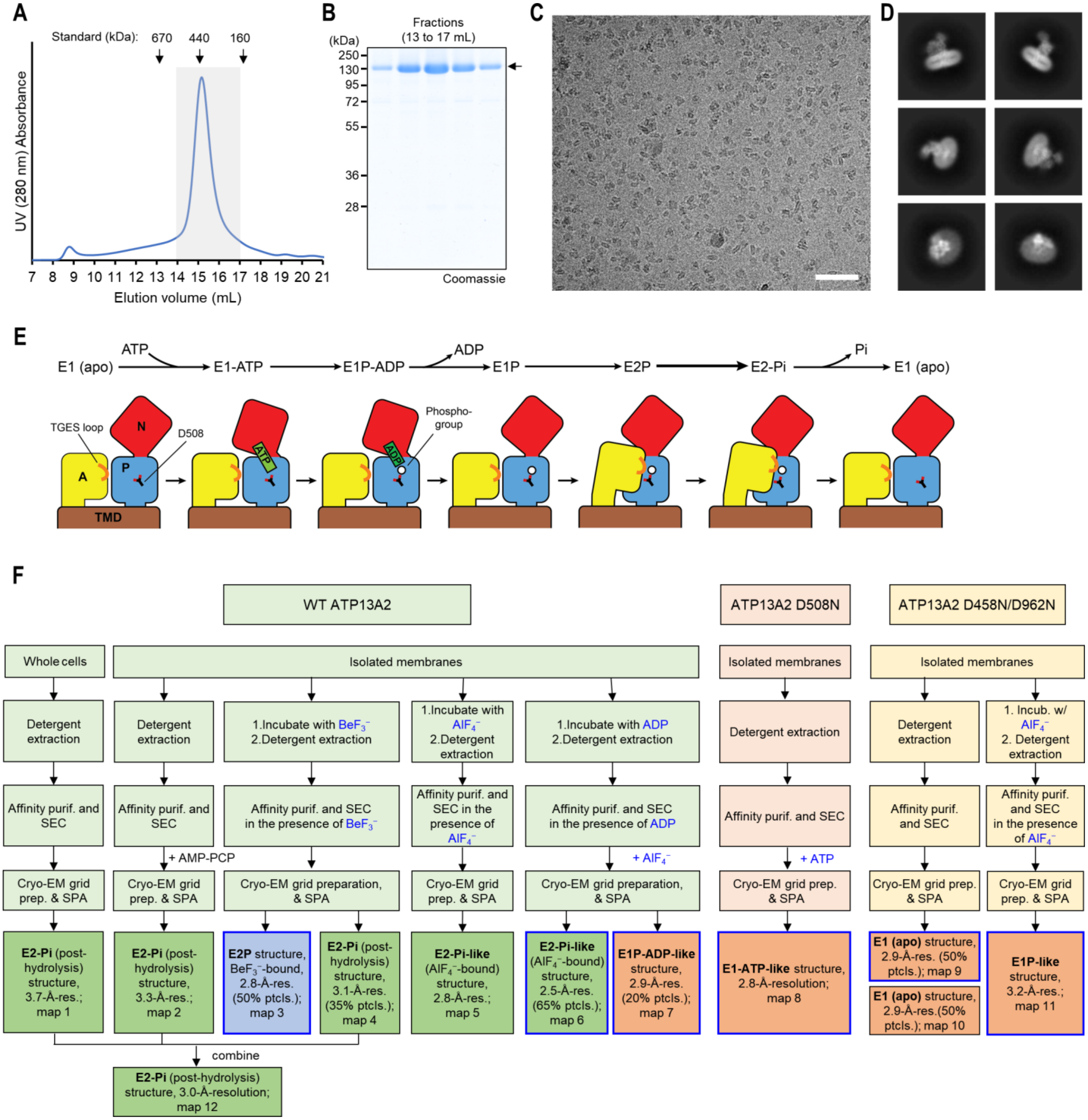


**Figure S1. Purification and cryo-EM analysis of human ATP13A2.**

(**A**) Superose-6 size-exclusion chromatography profile of affinity-purified ATP13A2. $AlF_4^-$-treated WT ATP13A2 is shown as a representative sample. The gray box indicates fractions used for cryo-EM analysis. (**B**) Coomassie-stained SDS gel of the Superose-6 fractions in A. (**C**) An example cryo-EM micrograph ($AlF_4^-$-bound WT ATP13A2). Scale bar, 50 nm. (**D**) Selected 2D class averages. The box width is 292 Å. (**E**) Schematic describing conformational changes of P-type ATPases in the Post-Albers cycle. Cytosolic domains are indicated by A (actuator), N (nucleotide-binding), and P (phosphorylation). The TGES ($Thr^{341}$-Gly-Glu-$Ser^{344}$) loop, which is responsible for catalyzing dephosphorylation, is also indicated (orange). (**F**) Flowchart summarizing the purification procedures and cryo-EM results. Blue-outlined maps are referenced as representatives of unique states of ATP13A2 in this study. Maps 1, 2, and 4 (and combined map 12) likely represent the product (post-hydrolysis) state of the dephosphorylation reaction. We also note that in map 2, nonhydrolyzable ATP analog β,γ-methyleneadonosine 5'-triphosphate (AMP-PCP) did not bind to ATP13A2. SPA, single particle analysis.

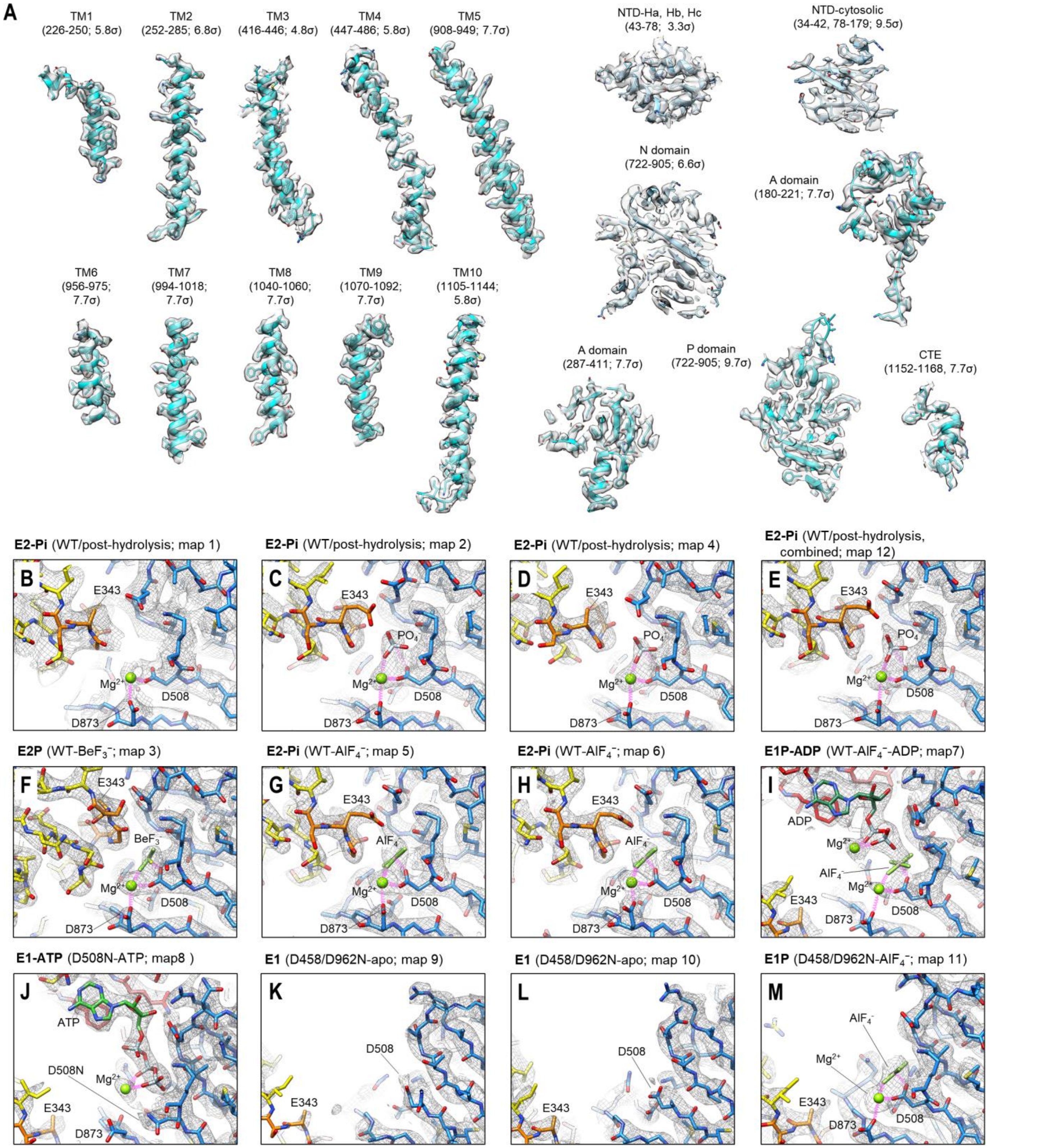


**Figure S2. Quality of the cryo-EM reconstructions and catalytic states.**

(**A**) Segmented EM densities and atomic models of ATP13A2. Shown are densities and models from the 2.5-Å-resolution $AlF_4^-$-bound structure (map 5), except for the NTD and N domain, which are from the 2.8-Å-resolution $BeF_3^-$-bound structure (map 3). (**B-M**) Cryo-EM density maps (mesh) and structures of the phosphorylation site in the different intermediate states of ATP13A2. Color scheme: yellow, A domain; orange, TGES loop; blue, P domain; red, N domain. We note that for some maps, the side chain of E343 (part of the TGES loop) was not modelled due to insufficient density features. In map 1 (panel B), the phosphate ion (Pi) density was not visible likely due to relatively lower (3.7 Å) resolution than the other E2-Pi maps (3.0–3.3-Å resolution).

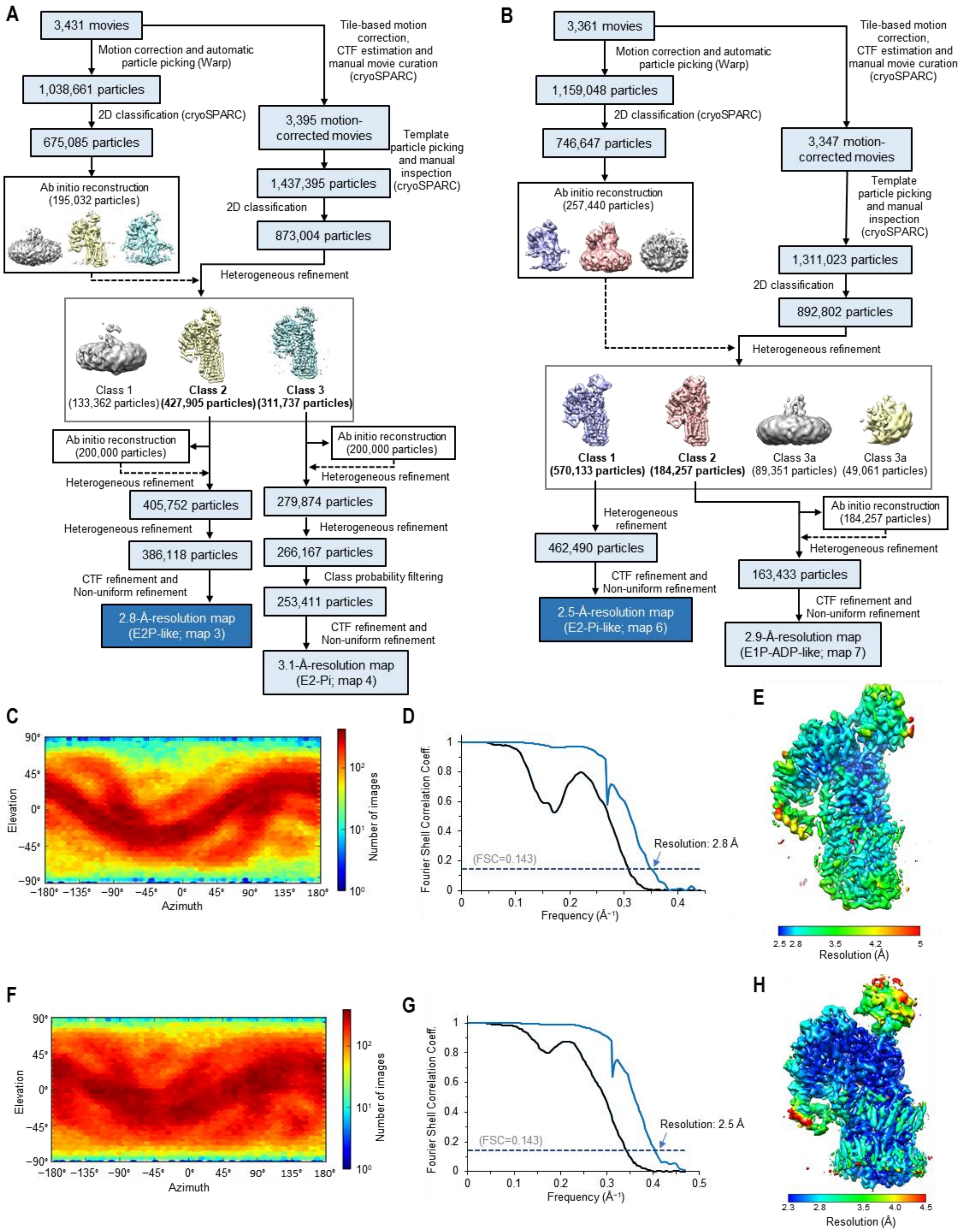


**Figure S3. Cryo-EM analysis of WT ATP13A2 prepared with $BeF_3^-$ or with ADP and $AlF_4^-$.**

**Figure S3. Cryo-EM analysis of WT ATP13A2 prepared with $BeF_3^-$ or with ADP and $AlF_4^-$.**

(**A**) Summary of single particle analysis for WT ATP13A2 prepared with $BeF_3^-$. (**B**) Summary of single particle analysis for WT ATP13A2 prepared with ADP and $AlF_4^-$. (**C**) Distribution of particle orientations in the final 3D reconstruction of the E2P-like structure (map 3). (**D**) Fourier shell correlation (FSC) curves between the two half maps of map 3. (**E**) Local resolution of map 3, indicated by a heat map. (**F**) Distribution of particle orientations in the final 3D reconstruction of map 6. (**G**) FSC between the two half maps of map 6. (**H**) Local resolution of map 6.

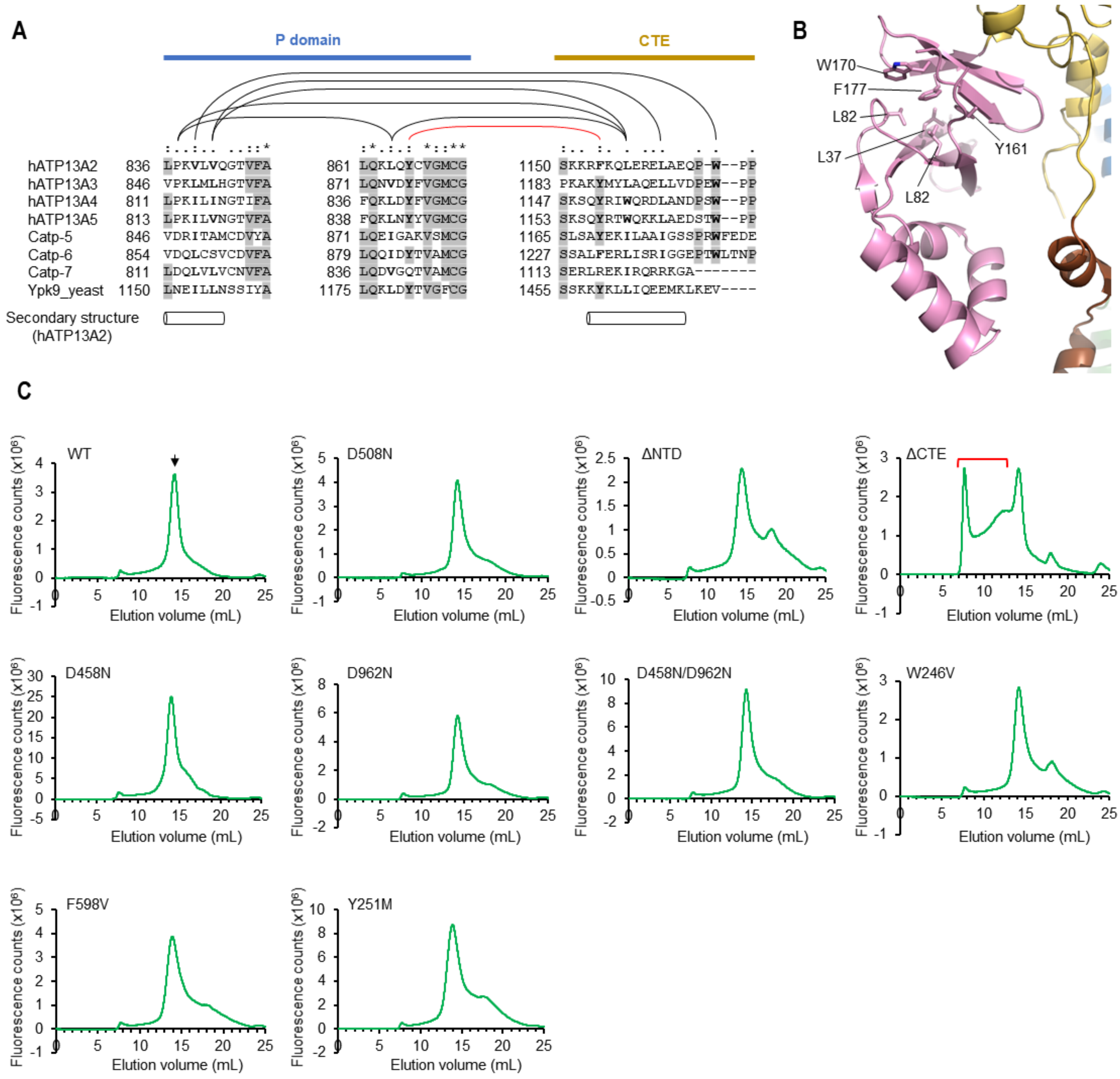


**Figure S4. Unique N- and C-terminal features of ATP13A2 and solution behaviors of ATP13A2 mutants.** (**A**) Sequence alignment between ATP13A2 homologs (P5B-ATPases) from human (ATP13A3-ATP13A5), *Caenorhabditis elegans* (Catp5-7), and *Saccharomyces cerevisiae* (Ypk9). Shown are the CTE region and parts of the P domain that interact with the CTE (for full sequences, see Supplemental item 1. Cylinders indicate position of alpha-helices, based on the ATP13A2 structure. Residues in gray shade are >50% identity. Lines indicate positions participating in hydrophobic interactions (black) and π-π stacking (red) in the ATP13A2 structures. Residues in bold are conserved aliphatic or aromatic amino acids for the interactions. (**B**) Structure of β-sandwich of the NTD. F177 (its mutation to valine is associated with hereditary early-onset Parkinson's disease called Kufor Rakeb Syndrome) and neighboring amino acids that form hydrophobic interactions are shown in a stick representation. (**C**) Fluorescence size-exclusion chromatography (FSEC) elution profiles of WT and mutant ATP13A2 after extraction of microsomes with DDM/CHS. A Superose 6 10/300 column was used. Black arrow, position of the main peak of WT ATP13A2. Red bracket, high molecular-weight aggregates.

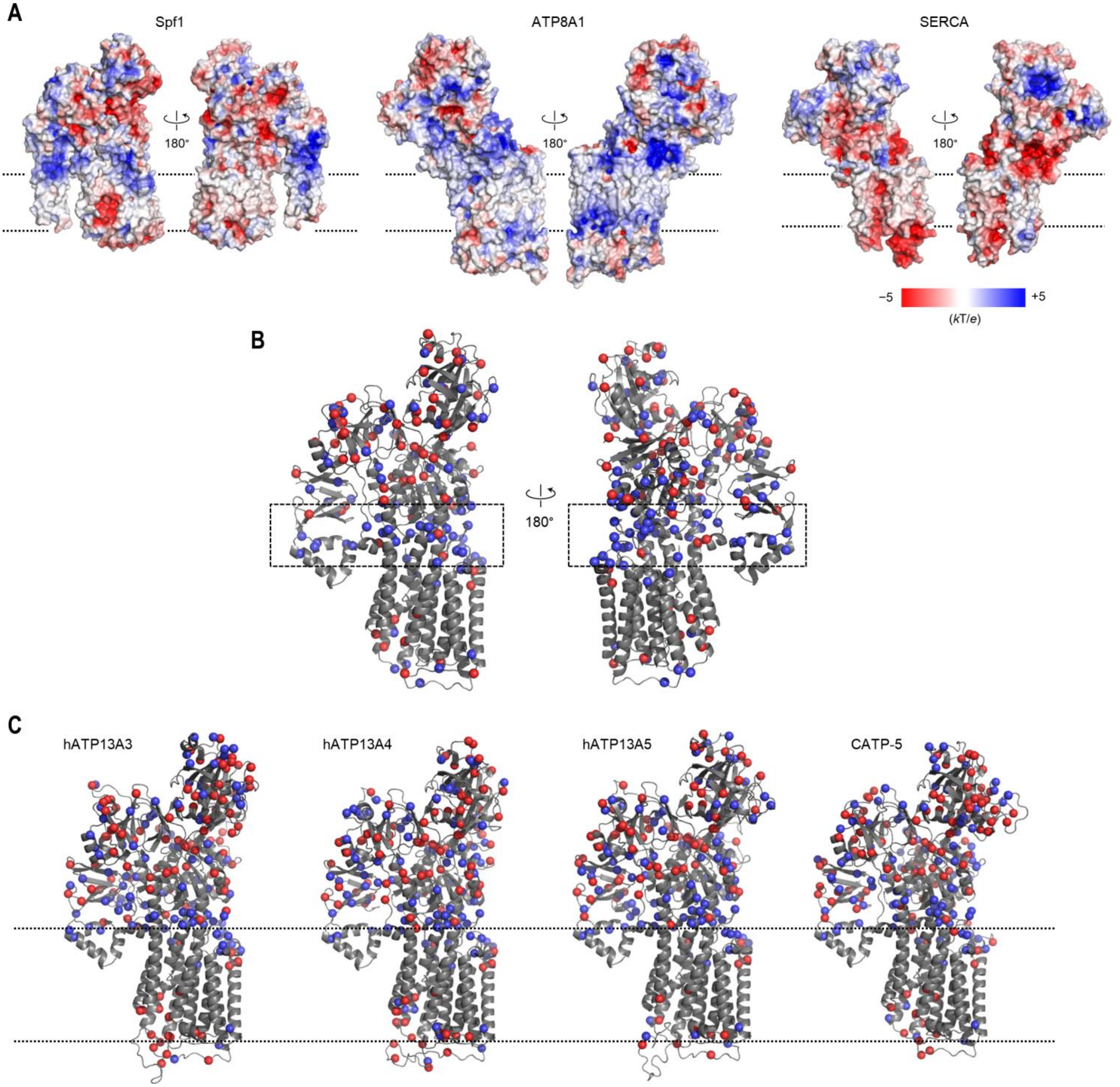


**Figure S5. Comparison of charge distributions among P-type ATPases.**
(**A**) As in Fig. 1D, but showing surface electrostatics of P5A-ATPase Spf1 (PDB: 6XMT), P4-ATPase ATP8A1 (PDB: 6K7M), and P2-ATPase SERCA (PDB: 3B9B). (**B**) Positions of basic (Arg and Lys; blue) and acidic (Asp and Glu; red) amino acids in ATP13A2 (the E2P structure). Dashed box indicates the region of the positively charged belt adjacent to the cytosol-membrane interface. (**C**) As in B, but with homology models of human P5B-ATPase isoforms (ATP13A3–ATP13A5) and the *C. elegans* ortholog (CATP-5). Dashed lines indicate the boundaries of the membrane.

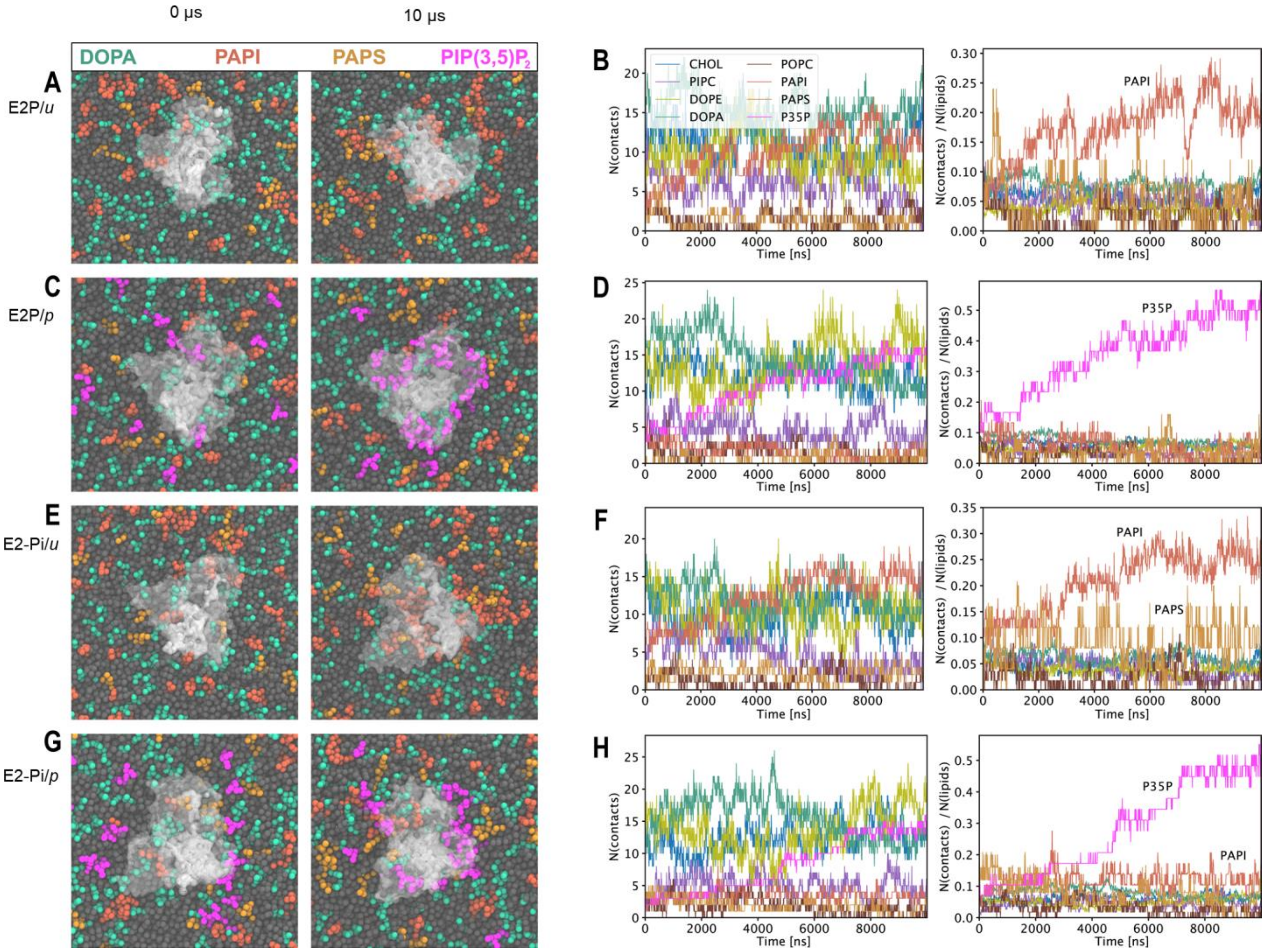


**Figure S6. Coarse-grained molecular dynamics (MD) simulations of ATP13A2 in model lysosomal membranes.**
(**A** and **B**) MD simulation with the E2P structure of ATP13A2 and a model membrane without PI(3,5)$P_2$. The panels in a show top-down views of ATP13A2 (white translucent surface) embedded in the membrane at the start of the simulation (left) and at 10μs simulation time (right). The panels in B show time-traces of protein-lipid contacts in the upper (cytosolic) leaflet over 10-μs simulation time (left panel, total number of contacts; right panel, contacts per lipid). The *u* membrane consists of 15.2% cholesterol (CHOL), 19.8% phosphatidylcholine (PIPC and POPC), 32.0% phosphatidylethanolamine (DOPE), 23.0% phosphatidic acid (DOPA), 3.0% phosphatidylserine (PAPS), and 7.0% phosphatidylinositol (PAPI) in the cytosolic leaflet, and 38.8% cholesterol, 30.8% PIPC, 15.4% POPC, and 15.0% sphingomyelin (DPSM) in the lumenal leaflet. In panels in A, negatively charged lipids are shown in colors as indicated (in bead representation) and other lipids are shown as gray beads. (**C** and **D**) As in A and B, but with 3.5% PI(3,5)$P_2$ (abbreviated as P35P) and PAPI reduced to 3.5% in the cytosolic leaflet of model membrane *p*. (**E**–**H**) As in A–D, but with the E2-Pi structure. Also see Movies S1–S4.

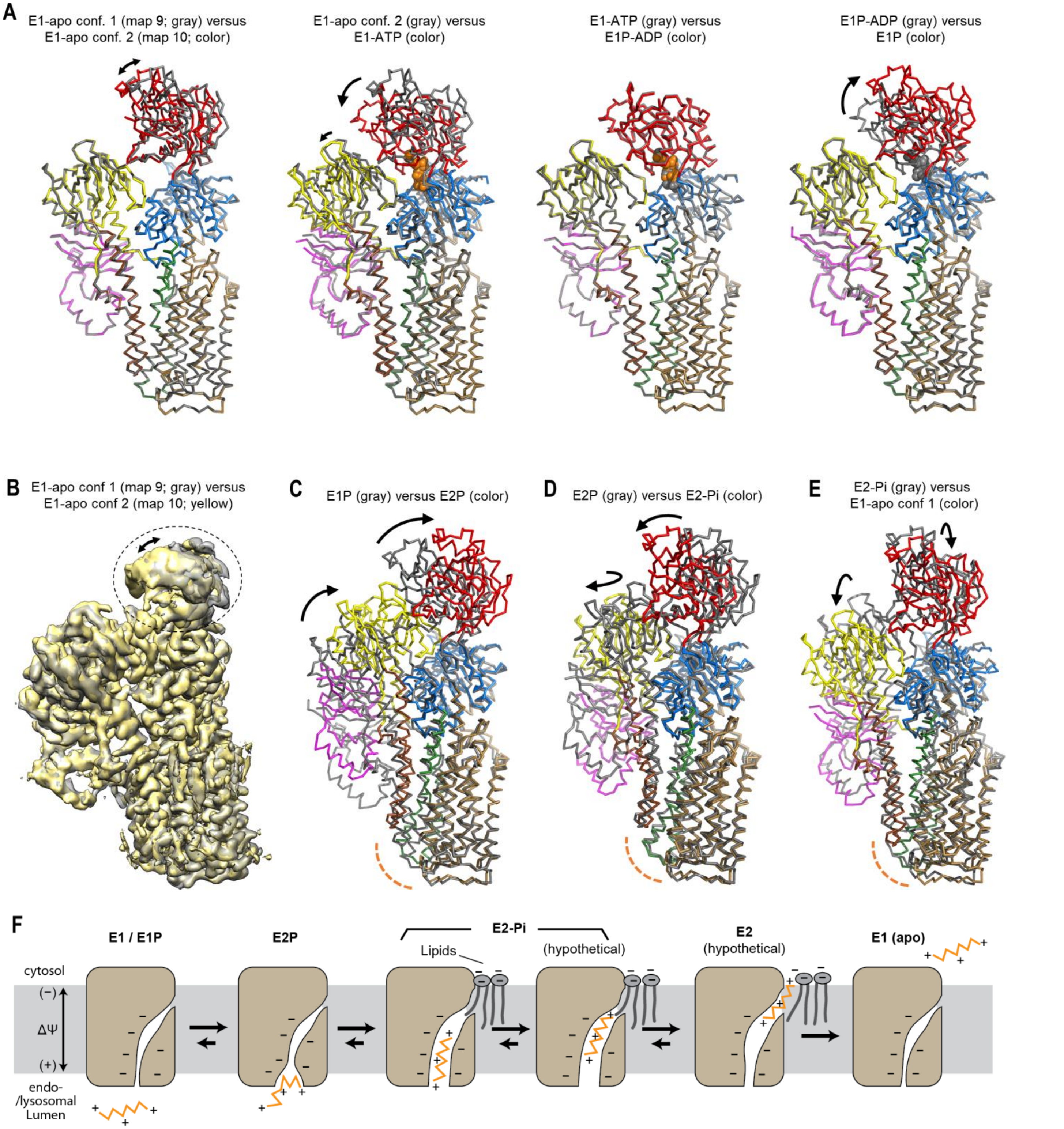


**Figure S7. Movements of the cytosolic domains and TMD during the catalytic cycle of ATP13A2.**
(**A**) Structural comparisons between different E1 intermediate states of ATP13A2, shown in a backbone-trace representation. The color scheme is the same as Fig. 1 A–C. Arrows indicate major domain movements between the compared structures. Nucleotides shown in a ball representation. Structures were aligned based on the P domain and TMs 5–10. (**B**) Comparison of the Cryo-EM density maps of two conformations observed for the E1 (apo) state (map 9 vs map 10) of the D458N/D962N mutant. Circle and arrows indicate the region (N domain) showing variability and motion, respectively. (**C**) As in A, but comparing E1P and E2P states. Orange dashed line indicates the location of the polyamine entrance. (**D**) As in C, but comparing E2P and E2-Pi states. (**E**) As in C, but comparing E2-Pi and E1 (apo) states. (**F**) Putative model for the polyamine-translocation mechanism of ATP13A2. Schematics for the TMD are shown. Late endosomal and lysosomal membranes are known to have a membrane potential ($\Delta\Psi$) of ~20 to 60 mV (positive in the lumen). We note that the exact protonation states of ATP13A2 (acidic amino acids in the substrate cavity) and the exiting polyamine molecule are unknown.

**Table S1. Additional cryo-EM data collection and refinement statistics**

| | WT<br>E2-Pi, post-hydrolysis<br>(map 1)<br>(EMD-xxxx) | WT<br>E2-Pi, post-hydrolysis<br>(map 2)<br>(EMD-xxxx) | WT<br>E2-Pi, post-hydrolysis<br>(map 4)<br>(EMD-xxxx) | WT<br>E2-Pi, $AlF_4^-$-bound<br>(map 5)<br>(EMD-xxxx) |
|---|---|---|---|---|
| **Data collection and processing** | | | | |
| Magnification | 64,000x | 64,000x | 64,000x | 64,000x |
| Voltage (kV) | 300 | 300 | 300 | 300 |
| Electron exposure ($e^-$/Å$^2$) | 50 | 50 | 50 | 50 |
| Defocus range (μm) | -1.0 to -2.5 | -0.7 to -2.4 | -0.7 to -2.5 | -0.7 to -2.4 |
| Pixel size (Å) | 0.911 | 0.911 | 0.911 | 0.911 |
| Symmetry imposed | *C*1 | *C*1 | *C*1 | *C*1 |
| Initial particle images (no.) | 605,982 | 1,001,323 | 1,437,395 | 1,716,479 |
| Final particle images (no.) | 164,606 | 173,886 | 253,411 | 593,369 |
| Map resolution (Å) | 3.7 | 3.3 | 3.1 | 2.8 |
| FSC threshold | 0.143 | 0.143 | 0.143 | 0.143 |
| Map resolution range (Å) | 3.1–10 | 2.8–9 | 2.7–18 | 2.4–12 |

**Table S2. Disease-associated missense mutations in ATP13A2**

| Mutation | Numbering in structure | Disease | Domain | Location in structure | Known protein defects | References |
|---|---|---|---|---|---|---|
| T12M | T12M | Early-onset Parkinson's disease (KRS) | NTD | Unknown | Decreased ATPase activity | (Di Fonzo et al., 2007; Podhajska et al., 2012; van Veen et al., 2020) |
| F182L | F177M | Early-onset Parkinson's disease (KRS) | NTD | Buried | Decreased protein stability, mislocalized to ER | (Estrada-Cuzcano et al., 2017; Ning et al., 2008; Podhajska et al., 2012; Wang et al., 2019) |
| A249V | A244V | Early-onset Parkinson's disease | TM1 | Lipid-facing | TBD | (Djarmati et al., 2009) |
| S282C | S277C | Early-onset Parkinson's disease | TM2 | Buried | TBD | (Djarmati et al., 2009) |
| I411M | I406M | Juvenile-onset amyotrophic lateral sclerosis (ALS) | A domain | Buried | TBD | (Spataro et al., 2019) |
| I441F | I436F | Early-onset Parkinson's disease (KRS) | TM3 | Lipid-facing | TBD, associated with A1069T | (Suleiman et al., 2018) |
| R449Q | R444Q | Early-onset Parkinson's disease | TM3 | Lumen-membrane interface | TBD | (Djarmati et al., 2009) |
| G504R | G499R | Early-onset Parkinson's disease (KRS) | P domain | Buried | Decreased protein stability, mislocalized to ER | (Di Fonzo et al., 2007; Podhajska et al., 2012; Wang et al., 2019) |
| T517I | T512I | Hereditary spastic paraplegia | P domain | Close to D508 | Loss of ATPase activity, mislocalized to ER | (Estrada-Cuzcano et al., 2017; van Veen et al., 2020) |
| G522V | G517V | Early-onset Parkinson's disease (KRS) | P–N connection | P–N connection | TBD | (Grunewald et al., 2012) |
| G533R | G528R | Early-onset Parkinson's disease (KRS) | N domain | Solvent exposed, beta turn | Decreased ATPase activity | (Di Fonzo et al., 2007; Estrada-Cuzcano et al., 2017; Podhajska et al., 2012; van Veen et al., 2020) |
| R709T | R704T | KRS, Hereditary spastic paraplegia | N domain | Partially solvent exposed | TBD | (Estiar et al., 2020; McNeil-Gauthier et al., 2019) |
| G720W | G715W | Hereditary spastic paraplegia | N domain | Buried | TBD | (Estiar et al., 2020) |
| A746T | A741T | Early-onset Parkinson's disease (KRS) | P domain | Buried | Decreased ATPase activity | (Podhajska et al., 2012; van Veen et al., 2020) |
| P842L | P837L | Early-onset Parkinson's disease (KRS) | P domain | CTE interface | TBD | (Abbas et al., 2017) |
| M854R | M849R | KRS, Neuronal ceroid lipofuscinosis | P domain | Buried | TBD | (Bras et al., 2012) |
| G877R | G872R | Early-onset Parkinson's disease (KRS) | P domain | Buried | Decreased protein stability, mislocalized to ER | (Podhajska et al., 2012; Santoro et al., 2011; van Veen et al., 2020) |
| G892D | G887D | Hereditary spastic paraplegia | P domain | Buried | TBD | (van de Warrenburg et al., 2016) |
| R980H | R975H | Early-onset Parkinson's disease | Loop TM6/7 | Cytosol-membrane interface | TBD | (Djarmati et al., 2009) |
| G1014S | G1009S | Early-onset Parkinson's disease | TM7 | Lipid-facing | TBD | (Chen et al., 2011) |
| L1059R | L1054R | Early-onset Parkinson's disease (KRS) | TM8 | Buried | Mislocalized to ER | (Park et al., 2011) |
| A1069T | A1064T | Early-onset Parkinson's disease (KRS) | Loop TM8/9 | Cytosol-membrane interface | TBD, associated with I441F | (Suleiman et al., 2018) |

KRS, Kufor-Rakeb syndrome. TBD, to be determined.
